# Synaptic plasticity in the orbitofrontal cortex explains how risk attitude adapts to the range of risk prospects

**DOI:** 10.1101/2020.09.08.287714

**Authors:** Jules Brochard, Jean Daunizeau

## Abstract

Many deleterious behaviors, such as procrastinating on urgent matters or sustaining a toxic relationship, are irrational. But is irrational behavior the incidental outcome of biological constraints imposed on neural information processing? In particular, can those constraints alter decisions, even when people know the consequences of alternative actions? Recent studies indicate that orbitofrontal neurons encode decision value in relative terms, i.e. value signals in OFC neurons are normalized with respect to the context. Value-based decisions may thus exhibit irrational context-dependence effects. A candidate explanation is “efficient coding”: OFC neurons may mitigate information loss by adapting their (bounded) output firing properties to the recent value range. This is seducing, because it suggests that relative value coding is the brain’s best attempt to produce rational behavior, given its own hard-wired biological constraints. However, whether the behavioral implications of this scenario are met, how it generalizes to realistic situations in which OFC neurons construct value from multiple decision-relevant attributes, and what its neurophysiological bases are, is unclear. Here, we address these issues by re-analyzing two open fMRI datasets from the OpenNeuro.org initiative, where people have to integrate prospective gains and losses to decide whether to gamble or not. First, we show that peoples’ risk attitudes critically depend on the range of gain prospects they are exposed to. Importantly, counter to simple efficient coding scenarios, differences in gain ranges induce progressive changes in peoples’ sensitivity to both gains and losses. Second, we use artificial neural network models to show that hebbian rewiring processes between attribute-specific and attribute-integration neurons predict (out-of-sample) both context-dependence effects in peoples’ risk attitude and multivariate patterns of fMRI activity in the OFC. Under mild conditions on distributed population codes for decision attributes, hebbian plasticity eventually translates the distribution of reweighted attribute signals towards the responsive range of integration neurons. In turn, integration units exhibit the known features of range adaptation, including (but not limited to), relative value coding. Our results demonstrate how hebbian plasticity within brain networks may result in range adaptation, eventually yielding complex though predictable irrational behavior.

## Introduction

Why do we maintain unrealistic expectations or engage in irresponsible conducts? Maybe one of the most substantial and ubiquitous violations of rationality is peoples’ sensitivity to modifications and/or manipulations of contextual factors that are irrelevant to the decision problem (Kahneman, 2011; Seymour and McClure, 2008). A prominent example is that peoples’ attitude towards risks depends upon whether alternative choice options are framed either in terms of losses or in terms of gains (Kahneman and Tversky, 2012). More generally, many forms of irrational behaviors stem from peoples’ relative (as opposed to absolute) perception of value, that is: value is perceived in relation to a contextual reference point. Because it provides a mechanistic interpretation of such relative/context-dependent decision processes, range adaptation in value-sensitive neurons is currently under intense scrutiny (Louie et al., 2013; Rangel and Clithero, 2012; Rigoli et al., 2016; Rustichini et al., 2017; Steverson et al., 2019). Neural range adaptation was first observed in the brain’s perceptual system: neurons in the retina normalize their response to incoming light in their receptive field w.r.t. to the illumination context, such that output firing rates span the variation range of surrounding light intensities (see Carandini and Heeger, 2012 for a review). Such divisive normalization effectively mitigates the information loss that would otherwise result from the limited firing range of light-sensitive neurons, hence implementing so-called *efficient coding* (Brenner et al., 2000; Laughlin, 1981; Valerio and Navarro, 2003; Wark et al., 2007). This idea was later extended to value coding, i.e. value-sensitive neurons were shown to normalize their response w.r.t. the set of alternative options within a given choice context and/or to the recent history of experienced/prospective rewards (Louie et al., 2013; Rangel and Clithero, 2012). This predicts how peoples’ preference should drift with the context, eventually yielding foreseeable forms of irrational behavior. In particular, temporal range adaptation effects have been the focus of intense research over the past decade, because they hold the promise of explaining persistent behavioral changes. In line with the existing literature on value processing in the brain, they have been repeatedly documented in non-human primates, mostly using electrophysiological recordings in the orbitofrontal cortex or OFC (Conen and Padoa-Schioppa, 2019; Kobayashi et al., 2010; Padoa-Schioppa, 2009; Tremblay and Schultz, 1999; Yamada et al., 2018), though similar effects have been demonstrated in the anterior cingulate cortex (Cai and Padoa-Schioppa, 2012) and the amygdala (Bermudez and Schultz, 2010; Saez et al., 2017). Although comparatively sparser, neural evidence for temporal value normalization in the human OFC and ventral striatum also exists (Burke et al., 2016; Cox and Kable, 2014; Elliott et al., 2008). Importantly, when included into computational models of value-based decision making, neural normalization and/or efficient coding partially explains apparent behavioral inconsistency away (Polanía et al., 2019; Zimmermann et al., 2018).

Having said this, the neurophysiological bases of range adaptation in value-sensitive neurons are virtually unknown and their behavioral consequences are debated (Khaw et al., 2017; Rustichini et al., 2017). For example, that overt preferences do not shift along with the observed changes in the value-sensitivity of OFC neurons is puzzling. A possibility is that range adaptation in OFC neurons may be “undone” by Hebbian-like plasticity mechanisms that fine-tune the synaptic efficacy of downstream “value comparison” neurons (Padoa-Schioppa and Rustichini, 2014; Rustichini et al., 2017). Implicit in this reasoning is the assumption that option values are typically considered as input signals to “value-sensitive” OFC neurons, which then transmit this information to downstream decision systems, in analogy to the transmission of light intensity information by “light-sensitive” neurons in the retina (Louie and Glimcher, 2012). But another possibility is that value coding in OFC neurons departs from the logic of efficient information transmission in the visual system (Burke et al., 2016; Conen and Padoa-Schioppa, 2019). For example, OFC neurons may be constructing (as opposed to receiving) value signals, out of input signals conveying information about possibly conflicting decision-relevant attributes (Lim et al., 2013; O’Doherty et al., 2021; Pessiglione and Daunizeau, 2021a; Raghuraman and Padoa-Schioppa, 2014). Under efficient coding, OFC neurons would not adapt to the range of (integrated) values; rather, they would adapt to the range of their attribute inputs. Although this would eventually alter OFC output value signals, the ensuing behavioral consequences would depend upon the nature of the neurophysiological mechanism that enable range adaptation (Louie et al., 2015).

Consider typical decisions under risk, which requires integrating attributes such as prospective gains and losses. Following early efficient coding models of perceptual systems, range adaptation may operate prior to attribute integration, at the level of neurons that process each attribute independently of each other (Soltani et al., 2012). This would predict that peoples’ sensitivity to each attribute would depend solely upon the recent history of the corresponding attribute. For example, if the range of prospective gains increases, then peoples’ sensitivity to gains should decrease, but their sensitivity to losses should remain constant. Alternatively, range adaptation may operate at the level of neurons that integrate attributes. For example, the synaptic gain of attribute neurons may undergo activity-dependant plasticity, eventually translating the distribution of reweighted attribute signals towards the responsive range of integration neurons. A candidate mechanism is Hebbian plasticity, which is a ubiquitous and ever-persistent mechanism that reinforces the synaptic strength between two neurons when they fire in synchrony, hence the “fire together, wire together” iconic Hebbian rule (Hebb, 1950). A plethora of electrophysiological studies have established its many variants, including, but not limited to, spike-timing dependent plasticity and long-term synaptic potentiation/depression (Fox and Stryker, 2017; Lisman, 2017; Shouval et al., 2010; Zenke and Gerstner, 2017). In the sensory domain, hebbian plasticity is the basic explanation behind the emergence of neuronal selectivity (Sadeh et al., 2015; Zhang et al., 1993). More precisely, hebbian plasticity determines the competition between synapses, so that neurons become unresponsive to some features while growing more responsive to others (Abbott and Nelson, 2000). When making decisions, such plastic competition of attribute inputs to OFC neurons is in fact necessary to construct value in a flexible and context-dependent manner (O’Doherty et al., 2021). Thus, although hebbian plasticity may *a priori* be seen as a hard-wired incidental constraint imposed on any neural system, it may in principle underlie range adaptation in OFC (and/or other value coding) neurons. In fact, under mild conditions regarding upstream population codes for decision attributes, the impact of hebbian plasticity is reminiscent of efficient coding, in the sense that integration neurons respond to their input history by increasing their output variability, yielding better resolved output value signals. However, this scenario differs from efficient coding on attribute-specific neurons, in that it likely implies between-attribute spill-over effects: e.g., changes in the range of gain prospects would likely alter peoples’ sensitivity to losses. In turn, one may reasonably ask whether hebbian plasticity may contribute to context-dependency effects in value-based decision making. This work is a first step in this direction.

In what follows, we focus on *risk attitude,* i.e. peoples’ tendency to accept or reject risky gambles given prospective gains and losses. We perform an entirely novel re-analysis of two independent fMRI datasets, which are made available in the context of the *openneuro.org* initiative (Poldrack et al., 2013). In both studies, participants are asked to accept or reject a series of gambles, but the two studies differ w.r.t. to range of prospective gains. First, we provide evidence that peoples’ risk attitude depends upon the gain context. Second, we show that the context-dependency of risk attitudes exhibits strong spill-over range adaptation effects, which precludes “pure” efficient coding at the level of attributes (i.e. gains and/or losses). Third, we ask whether such irrational behavioral changes may be explained by range adaptation at the level of integration neurons (induced by hebbian plasticity). At this point, we note that quantifying the neural and behavioral implications of this scenario is not trivial, which is why we resort to artificial neural network (ANN) modelling. We consider an idealized “value constructing” network, where value is read out of a layer of integration neurons, which receive inputs from upstream layers of attribute-specific neurons (O’Doherty et al., 2021; Pessiglione and Daunizeau, 2021b). We test whether, when equipped with hebbian plasticity, these ANNs provide predictions about peoples’ trial-by-trial choices that generalize (out-of-sample) across risk range contexts. We then compare the activity patterns of artificial integration units to multivariate fMRI patterns, using a variant of representational similarity analysis or RSA (Diedrichsen and Kriegeskorte, 2017; Diedrichsen et al., 2020; Kriegeskorte, 2008). All relevant computational and statistical details are summarized in the Methods section.

## Results

We now present our re-analysis of the NARPS dataset (Botvinik-nezer et al., 2019). This dataset includes two studies, each of which is composed of a group of 54 participants who make a series of decisions under risk. On each trial, a gamble was presented, entailing a 50/50 chance of gaining an amount G of money or losing an amount L. As in Tom et al. (2007), participants were asked to evaluate whether or not they would like to accept or reject the gambles presented to them. In the first study (hereafter referred to as the “narrow range” group), participants decided on gambles made of gain and loss levels that were sampled from the same range (G and L independently varied between 5$ and 20$). In the second study (hereafter: the “wide range” group), gain levels scaled to double the loss levels (L varied between 5$ and 20$, and G independently varied between 10$ and 40$). Importantly, both groups experience the exact same range of losses. In both studies, all 256 possible combinations of gains and losses were presented across trials (see Methods section). Importantly, the gambles’ outcomes were not revealed until the end of the experiment.

First, we ask whether peoples’ attitude towards risks exhibits range adaptation. In our context, range adaptation implies that the variation in peoples’ tendency to accept risky gambles should approximately span the range of gambles’ values that they are exposed to. In Figure 1 below, the upper-left panel shows the observed probability of gamble acceptance as a function of gambles’ expected value (i.e. EV=(G-L)/2), for both groups.

**Figure 1:**
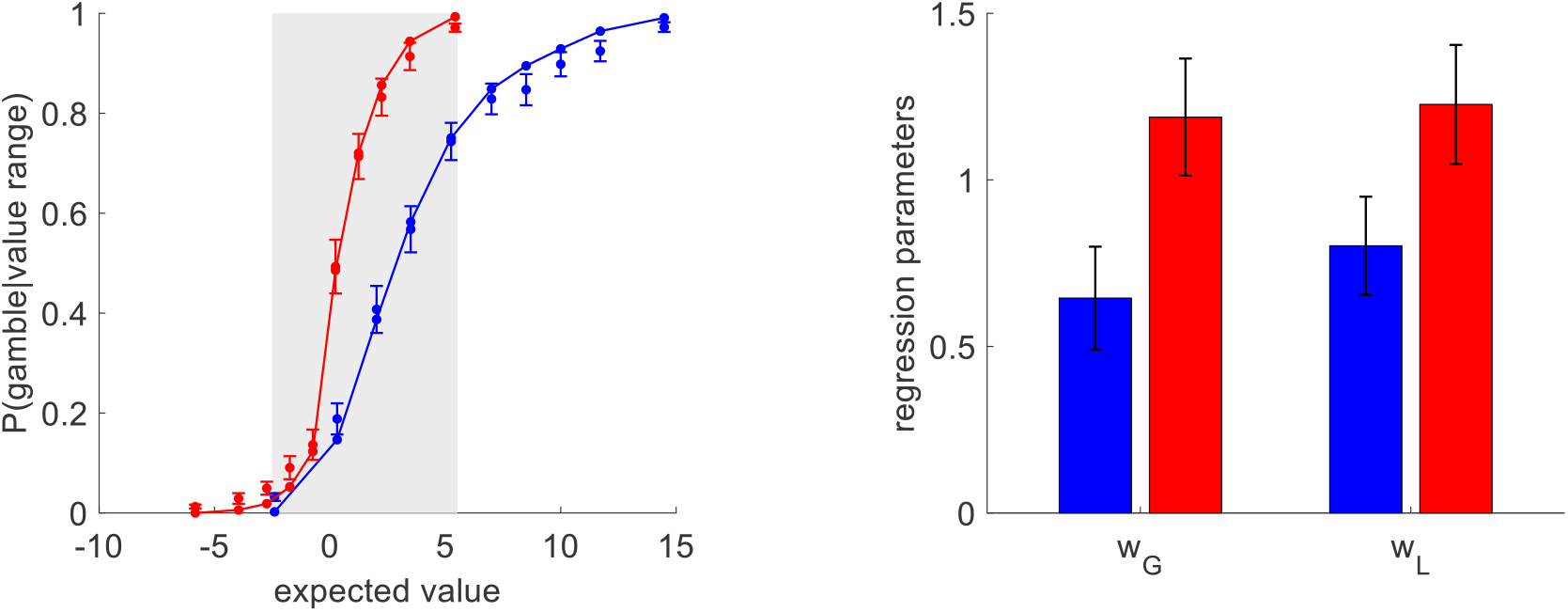
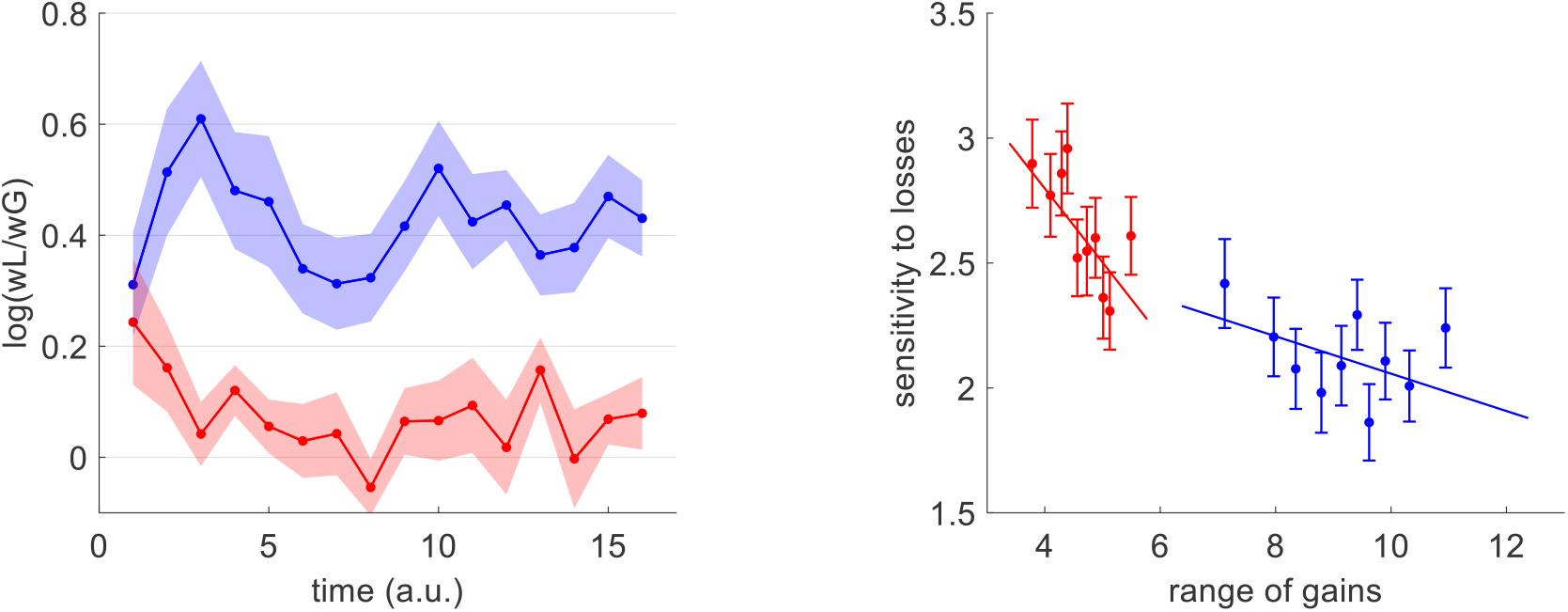
Choice inconsistency and range adaptation. **Upper-left panel:** the group-average probability of gamble acceptance (y-axis) is plotted against deciles of gambles’ expected value (x-axis), in both groups (red: narrow range, blue: wide range). Dots show raw data (errorbars depict s.em.), and plain lines show predicted data under a logistic regression model (see main text). The grey shaded area highlights the range of expected values that is common to both groups. **Upperright panel:** Estimates of sensitivity to gains (wG) and losses (wL) for both groups, under the logistic model (same color code as upper-left panel, errorbars depict s.e.m.). **Lower-left panel:** Temporal dynamics of group-average loss aversion (y-axis, same color code) is plotted against time (a.u., x-axis). Shaded areas depict s.e.m. **Lower-left panel:** Sensitivity to losses (y-axis, same color code) is plotted against deciles of range of gains (x-axis). Errorbars show s.e.m. and plain lines depict regression lines.

Overall, the average gambling rate of people from the wide range group (65% ±2%) is much higher than that of people from the narrow range group (44% ±2%), and the group difference is significant (p<10-4, F=44.8, dof=101). This is of course expected, given that people from the wide range group are exposed to gambles with higher value on average. However, and most importantly, within the range of expected values that is common to both groups, people from the wide range group are *less* likely to gamble than people from the narrow range group. Here, the average gambling rate of people from the wide range group is 41% ±3%, whereas it is 54% ±3% for people in the narrow range group, and this group difference is significant (p=0.003, F=9.2). This difference can only be due to the context in which people made these decisions, which is more favorable (higher gain prospects on average) in the wide range group. This is the hallmark of range adaptation.

To investigate this effect, we first performed a within-subject logistic regression of trial-by-trial choice data onto gains and losses (including an intercept, see Methods section). This regression accurately explains 91% ± 0.7% (resp. 87% ± 0.8%) of individual choices in the narrow (resp. wide) range group (cf. plain lines in upper-left panel of Figure 2). A random effect analysis on regression parameters failed to identify a difference in the constant gambling bias (mean difference of intercept estimates: p=0.82, F=0.05). However, peoples’ sensitivity to both gains and losses are significantly higher in the narrow range group than in the wide range group (w_G_: p<10^-4^, F=29.3; w_L_: p=0.001, F=11.1 ; cf. upper-right panel of Figure 1). This means that manipulating the range of gain prospects changes peoples’ sensitivity to both gains and losses. But this change seems to be slightly asymmetric, i.e. the gain range context does not alter peoples’ sensitivity to gain and loss prospects in the same way. To quantify this, we derived indices of individual loss aversion, which we define as the log-transformed ratio of loss sensitivity to gain sensitivity, i.e. log(w_L_/w_G_) (Tom et al., 2007). We found that people from the wide range group exhibit significant loss aversion (mean loss aversion index=0.41, sem=0.05, p<10^-4^) whereas people from the narrow range group do not (mean loss aversion index=0.037, sem=0.05, p=0.49), and the ensuing group difference is significant (p<10^-4^, F=24.3). This difference in loss aversion explains the observed difference in peoples’ attitude towards risks within the common range of gambles’ expected values.

**Figure 2:**
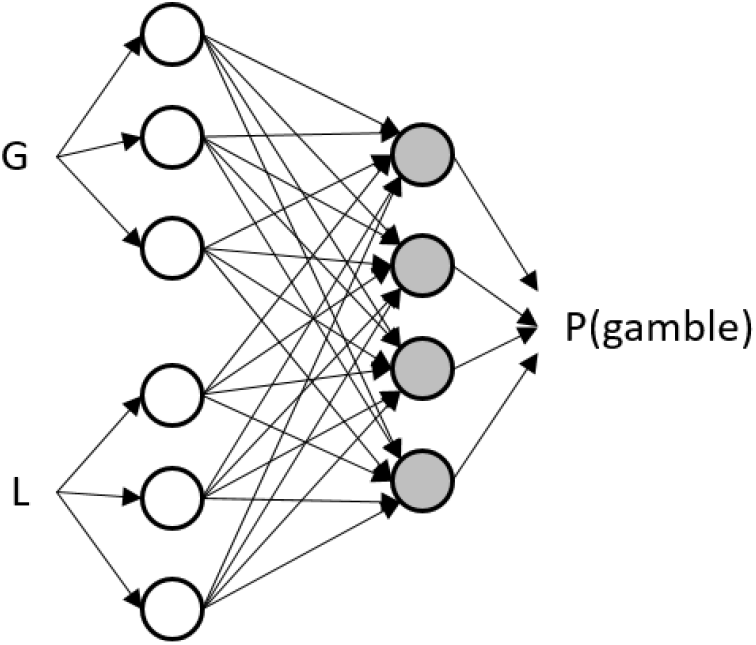
Schematic structure of the ANN. Trial-by-trial gains (G) and losses (L) first enter attribute-specific units, which then send their outputs to integration units (grey circles). The outputs of these units are then combined to yield trial-by-trial gamble acceptance decisions, using a linear decoder passed through a sigmoid mapping. When equipped with hebbian plasticity, the connections weights that relate attribute-specific units to integration units are modified in proportion to the amount of instantaneous covariance between their output firing rates.

But how did this loss aversion difference eventually ensue? To address this question, we repeated the within-subject logistic regression, this time on consecutive chunks of 16 trials (see Methods section). The resulting temporal dynamics of loss aversion are shown on the lower-left panel of Figure 1. At the start of the experiment (first 16 trials), loss aversion is significant in both groups (narrow range: mean loss aversion index=0.38, sem=0.1, p<10^-4^, wide range: mean loss aversion=0.24, sem=0.1, p=0.02), and there is no significant difference between groups (p=0.33, t=0.45). However, as people are exposed to more gambles, loss aversion in both groups tends to spread apart: the difference between groups starts becoming significant after 32 trials (p=0.008, t=2.43) and stays significant thereafter (all p<0.01). We then wondered whether this behavioral range adaptation effect can be explained under the efficient coding hypothesis. Recall under its simplest variant, efficient coding would operate on neurons that transmit information regarding either gains or losses, which would then adapt their sensitivity to their respective input range. First, we reasoned that, under efficient coding, temporal changes in peoples’ sensitivity to gains and losses should be driven by temporal changes in gain and/or loss ranges. For example, if the past few gambles involved a small range of gains, then peoples’ sensitivity to gain should increase. We thus regressed peoples’ sensitivity to gains and losses against the range of gains and losses (and their interaction), across temporal windows. We found that peoples’ sensitivity to gains negatively correlates with gain ranges within the narrow range group (p=0.02, F=5.43, df=762), but this was not the case within the wide range group (p=0.51, F=0.42, df=817). We also found that, in both groups, peoples’ sensitivity to losses negatively correlates with gain ranges (narrow range group: p=0.001, F=10.5, df=762 ; wide range group: p=0.045, F=4.04, df=817 ; see lower-right panel of Figure 1 and Supplementary Material). This is important, because efficient coding on neurons that transmit loss information cannot explain such between-attribute spill-over effect. Second, we reasoned that, under efficient coding, those brain regions that respond to gains and/or losses in both groups, should also show a between-group difference in sensitivity to gains and/or losses. Using fMRI univariate analyses (FWE corrected, see Methods), we found many brain regions that exhibited a significant group difference in sensitivity to either gains or losses, including the bilateral anterior insula (left: p_FWE_=0.053, right: p_FWE_=0.023), the ACC (p_FWE_=0.013), the right DLPFC (pFWE=0.001) and the right posterior parietal cortex (p_FWE_<10^-4^). However, hemodynamic activity in these regions does not respond significantly to either gains or losses. Alternatively, those regions that respond significantly to gains and/or losses in both groups (including, e.g., the vmPFC: p_FWE_<10^-4^ and the left OFC: p_FWE_=0.014) do not show a significant group difference in gain/loss sensitivity. In brief, when testing for the conjunction of either a gain or a loss effect in both groups, and a difference of this effect across groups, nothing in the brain ever reaches statistical significance (even with a more lenient significance threshold).

Taken together, those results dismiss simplistic efficient coding scenarios that would operate on neurons that transmit gain/loss information. But another possibility is that range adaptation operates on neurons that integrate information about prospective gains and losses to construct the subjective value of gambles. Qualitatively, range adaptation may result from hebbian rewiring processes that can translate the distribution of reweighted gain and loss signals towards the responsive range of integration neurons. It is, however, difficult to derive qualitative predictions from this type of scenario. Thus, we approach this problem using quantitative ANN models, equipped with hebbian plasticity mechanisms. We considered ANNs composed of two layers (see Methods): (i) an attribute-specific layer further divided into two sets of units that differ in terms of their inputs (either trial-by-trial gains or losses), and (ii) an integration layer receiving outputs from both attribute-specific units (see Figure 2 below).

By assumption, the gamble’s value is read out from the response pattern in the integration layer using a linear population code (Pessiglione and Daunizeau, 2021a). Note that, without a mechanism that modifies the units’ response, such ANNs cannot exhibit range adaptation. These ‘static’ ANNs will thus serve as reference points for our analyses. Following our behavioral results above, we modelled range adaptation effects at the level of the integration layer. In brief, hebbian plasticity strengthens (resp. weakens) pairwise synaptic connections between attribute and integration units in proportion to the covariance of their activity. Two global parameters control this “fire together – wire together” rule: the threshold parameter (λ_HP_) sets the amount of covariance above which connections increase (as opposed to decrease), and the rate parameter (α_HP_) controls the speed at which Hebbian plasticity modifies pairwise synaptic weights. Note that the net effect of hebbian plasticity may depend upon the shape of the units’ activation function (i.e. the relationship between their input’s magnitude and their output response). We thus equipped neuronal units with either monotonic (sigmoidal) or bell-shaped (pseudo-gaussian) activation functions. Each participant’s trial-by-trial choice sequence data was fitted with two candidate ANNs (static ANN, and ANN with hebbian plasticity i.e. HP-ANN), under either monotonic or bell-shaped units’ activation functions. In what follows, we refer to the models’ predictions about fitted behavioral data as models’ *postdictions.* We note that empirical distributions of ANN parameter estimates are reported in the Supplementary Materials. We also performed counterfactual model simulations: for each subject and each fitted model, we simulated the trial-by-trial gamble acceptances that would have been observed, had this subject/model be exposed to the sequence of prospective gains and losses that each subject *of the other group* was exposed to(see Methods). These out-of-sample predictions provide a strong test of each model’s generalization ability. Figure 3 below shows both postdictions and out-of-sample predictions of the four variants of ANN models (static versus plastic, with sigmoid versus gaussian activation functions).

**Figure 3:**
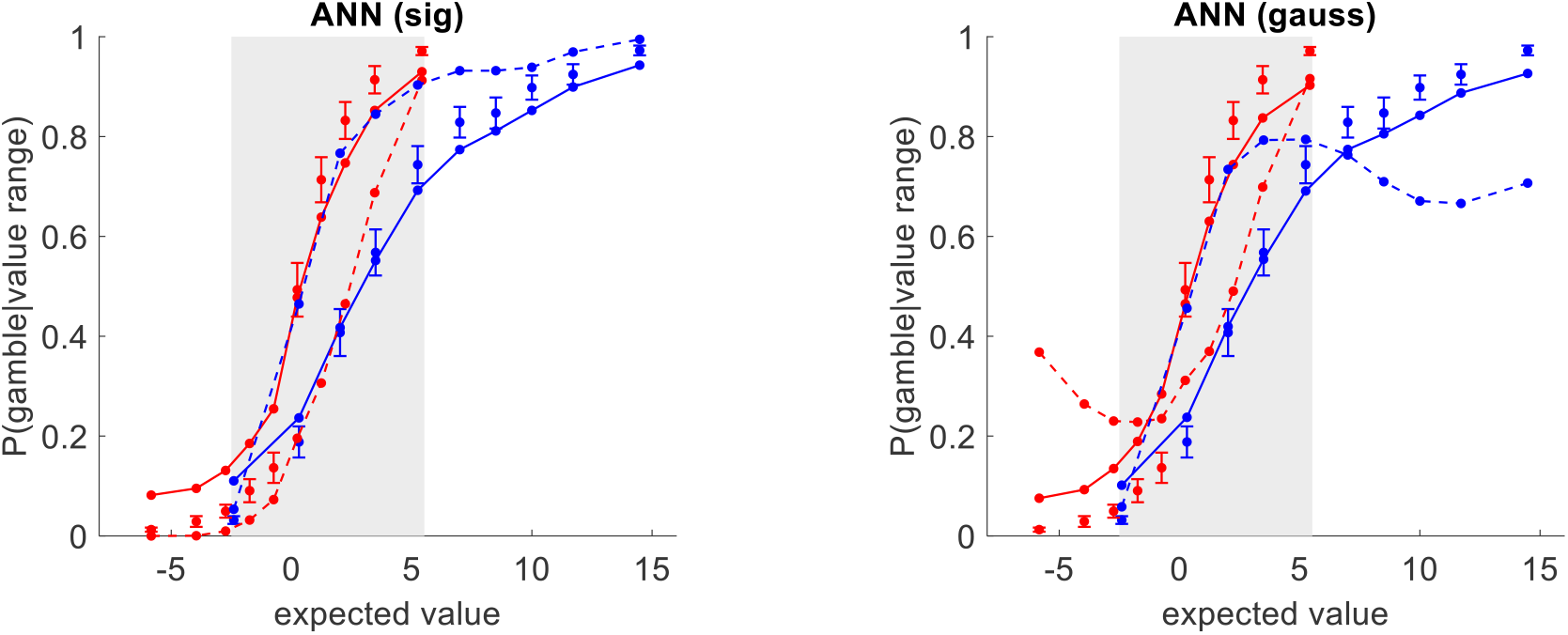
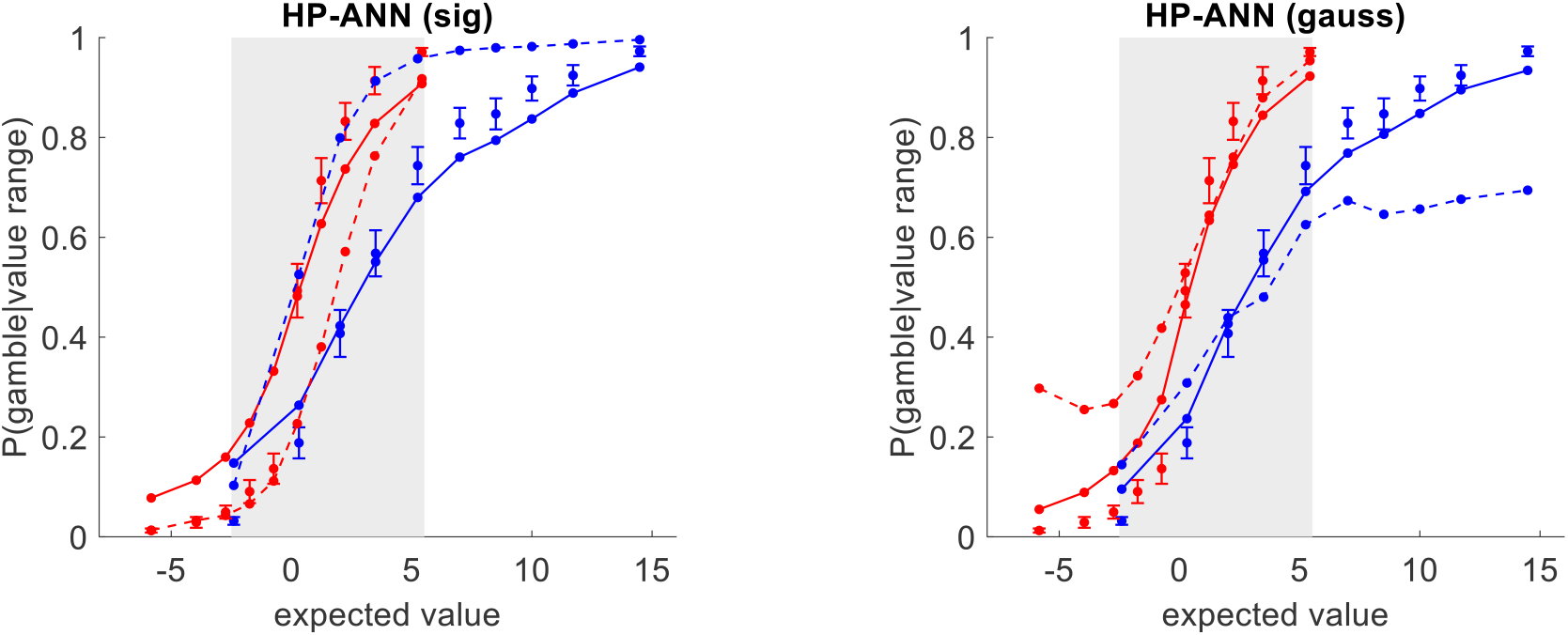
ANN analysis of behavioral data: out-of-sample predictions. **Upper-left panel:** The probability of gamble acceptance under static ANNs with sigmoid activation functions (y-axis) is plotted against gambles’ expected value (x-axis), in both groups (red: narrow range, blue: wide range). Dots show raw data (errorbars depict s.em.). Plain and dashed lines show postdictions and out-of-sample predictions (see main text), respectively. The grey shaded area highlights the range of expected values that is common to both groups. **Upper-right panel:** Same as upper-left panel, with gaussian activation functions. **Lower-left panel:** Probability of gamble acceptance under ANN with *Hebbian plasticity* (HP-ANNs) with sigmoid activation functions (same format as upper-right panel). **Lower-right panel:** Same as lower-left panel, with gaussian activation functions.

First, all model *postdictions* accurately describe the qualitative group difference in risk attitudes. In addition, all candidate ANNs perform much better than the logistic model, i.e. they all yield a significantly higher percentage of explained variance (all p<10^-5^, mean R^2^ of about 84%; see Table S3 of the Supplementary Material). Note that most of the fit improvement lies around the indifference point, where gains and losses balance out (see Figure S6 in the Supplementary Material). But did models capture a mechanism that faithfully generalizes to different gain contexts (i.e., across groups)? First, static ANNs do not yield accurate out-of-sample predictions. This is expected, because static ANNs cannot exhibit range adaptation. Thus, they tend to leave risk attitude unchanged within the common range of expected values, and simply extrapolate postdicted behavior outside that range (as for the logistic model, see Figure S1 in the Supplementary Material). In other terms, within the range of expected values that is common to both groups, static ANNs wrongly predict that risk aversion should be lower in the wide range group than in the narrow range group (mean gambling rate group difference=-33%, p<10^-4^, F=54.3). The situation is quite different for HP-ANNs with Gaussian activation functions, which yield rather accurate out-of-sample predictions of peoples’ risk attitude within the common range of expected values. In particular, those HP-ANNs correctly predict that risk aversion should be higher in the wide range group than in the narrow range group (mean gambling rate group difference=18%, p<10^-4^, F=13.6). This suggests that HP-ANNs (with gaussian activation functions) capture the essence of the range adaptation mechanism that underlies the context-dependency of peoples’ risk attitude.

We now aim at evaluating the neural evidence for or against candidate ANNs. We reasoned that if range adaptation in OFC neurons mediates the effect of the range of gain prospects on peoples’ risk attitudes, then within-subject fMRI activity patterns in the OFC should resemble corresponding within-subject activity patterns in ANNs’ integration layers in both groups of participants. We approach this problem using representational similarity analysis (RSA). This allows us to compare the trial-by-trial multivariate activity patterns of candidate ANNs with those of fMRI signals in the OFC, without any additional ANN parameter adjustment. In brief, we compute within-subject Representational Dissimilarity Matrices or RDMs, which measure the distance between activity patterns for each pair of trials. We then test for the correlation between ANN-based and fMRI-based RDMs. Here, we consider three subregions of the OFC: left/right lateral OFC (i.e. Brodmann areas 13), and medial OFC (i.e. vmPFC). Figure 4 below summarizes the RSA results, for HP-ANNs and static ANNs (with gaussian activation functions), in terms of the group-average RDM correlations (see Methods).

**Figure 4:**
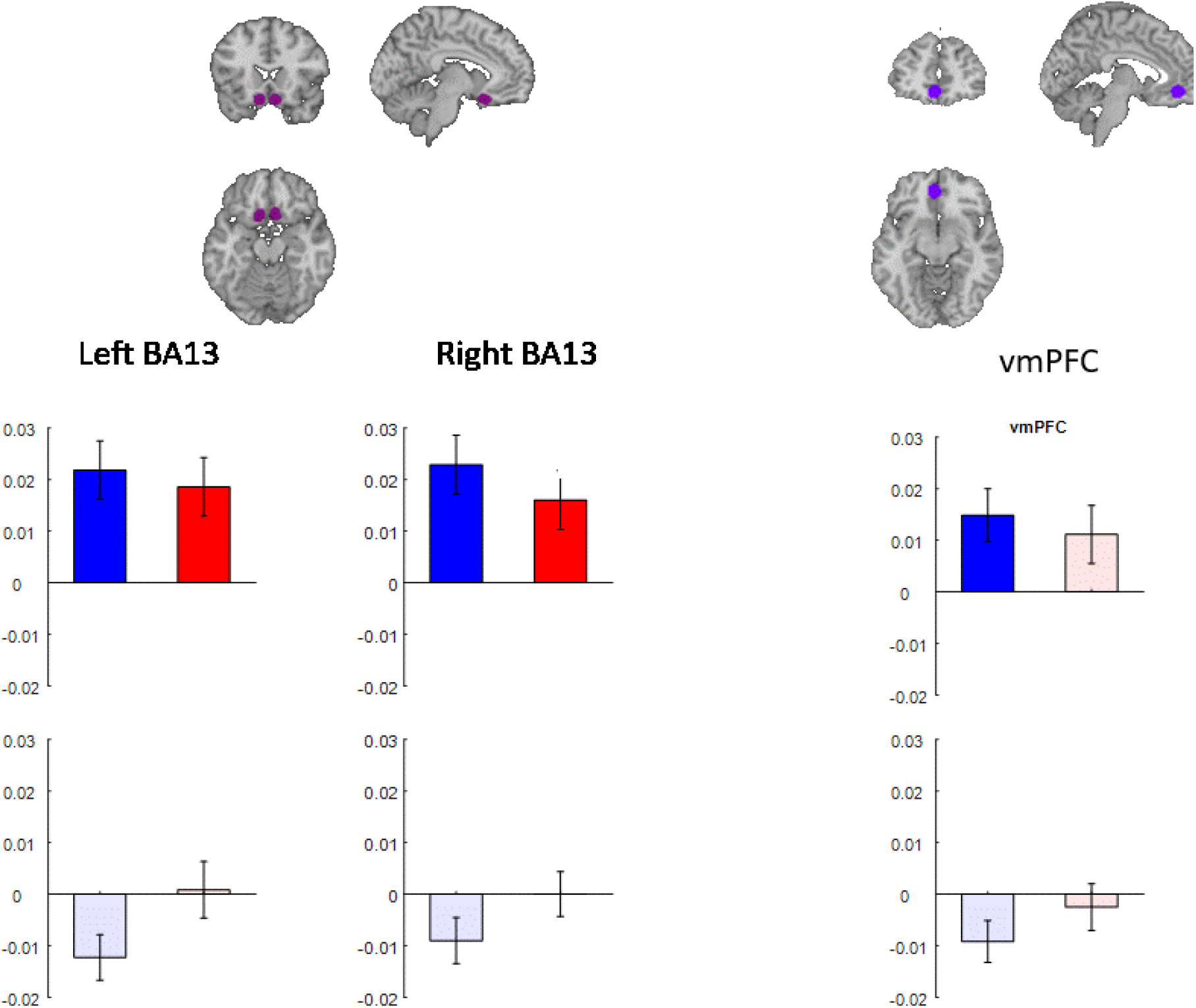
ANN analysis of fMRI data: RSA results. Each column shows the RSA results of a given OFC subregion (from left to right: left BA13, right BA13 and vmPFC). Each panel shows the group average RDM correlation ρ for HP-ANNs (upper row) and static ANNs (lower row), for each group of participants (blue: wide range group, red: narrow range group ; errorbars depict s.e.m). Plain bars depict statistical significance (at FPR=5%, Bonferroni correction across OFC subregions), whereas shaded bars show non significant results.

First, no static ANN reaches statistical significance, in any OFC subregion (all p>0.07, uncorrected). Second, HP-ANNs reach statistical significance in all OFC subregions, in each group of participant (all p<0.0035, uncorrected), except for vmPFC in the narrow range group (p=0.027, uncorrected). When comparing ANN models with each other using an RDM-based model comparison approach (see Supplementary Material), we found that HP-ANNs exhibit significantly higher RDM correlations than other ANNs in the wide range group (right BA13: p=0.009, left BA13: p=0.01, vmPFC: 0.03), though evidence is weaker in the narrow range group (right BA13: p=0.12, left BA13: p=0.18, vmPFC: p=0.2).

For completeness, we also performed the same RSA analysis in other regions of interest of the prefrontal cortex (ACC and bilateral DLPFC) as well as in the bilateral ventral striatum, bilateral amygdala, bilateral insula and PCC. Note that the OFC results are left qualitatively unchanged when correcting for multiple comparisons across all these regions of interest (as opposed to just left/right BA13 and vmPFC). Also, none of these regions show a significant RDM correlation with static ANNs. Having said this, in addition to bilateral BA13, RDM correlations of HP-ANNs reach statistical significance (in both groups) in bilateral DLPFC (p<0.001), bilateral ventral striatum (p<0.0007) and right amygdala (p<0.0009). We refer the interested reader to the Supplementary Material.

Taken together these results provide converging evidence that Hebbian plasticity is a likely explanation for range adaptation of risk attitude. But do HP-ANNs exhibit all the critical features of range adaptation? To address this question, we analyzed HP-ANNs using the same regression analyses than those we used for participants’ behavior (e.g., we performed logistic regressions with and without sliding temporal windows, see Methods). We then compared the results of these analyses to participants’ data. Figure 5 below summarizes the corresponding posterior checks.

**Figure 5:**
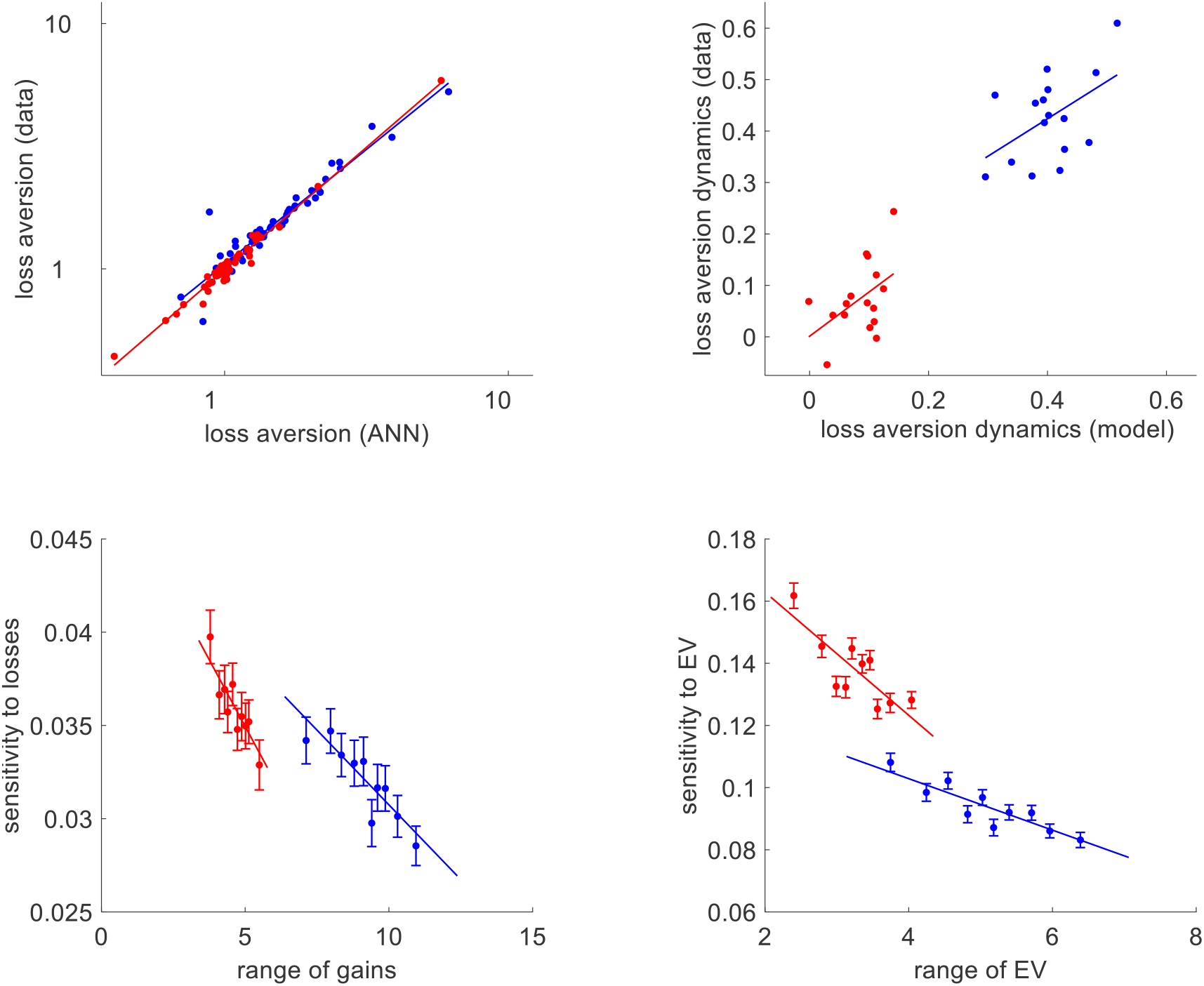
HP-HNNs (with Gaussian activation functions). **Upper-left panel**: Observed/raw loss aversion (y-data) is plotted against HP-ANN’s postdicted loss aversion (x-data). Each dot is a subject (red: narrow range, blue: wide range), and plain lines depict linear regression lines. **Upper-right panel:** Mean loss aversion dynamics (y-axis) is plotted against HP-ANN’s postdicted loss aversion dynamics (x-axis). Each dot is a temporal window (same color code). **Lower-left panel**: the sign-rectified sensitivity of HP-ANNs’ integration units to prospective losses (y-axis) is plotted against bins of prospective gain ranges (x-axis). **Lower-right panel**: the rectified sensitivity of HP-ANNs’ integration units to expected value (y-axis) is plotted against bins of expected values (x-axis).

First, we reasoned that, owing to the nonlinearity of range adaptation in integration neurons, within-subject sequences of gain and loss prospects may partially drive (context-dependent) inter-individual differences in loss aversion. Thus, a minimal requirement is that HP-ANNs accurately postdict inter-individual differences in loss aversion. Indeed, we find that the correlation between observed and postdicted loss aversion index (across subjects) is significant in both groups (wide range group: r=0.95, p<10^-4^; narrow range group: r=0.98, p<10^-4^; see upper-left panel of Figure 5). Second, we asked whether HP-ANNs accurately capture the progressive changes of loss aversion that is induced by the repeated exposure to either wide or narrow gain ranges. We find that the correlation between observed and postdicted temporal dynamics of loss aversion (across temporal windows) is significant in both groups (wide range group: r=0.50, p=0.024; narrow range group: r=0.46, p=0.035, see upper-right panel of Figure 5).

Third, we asked whether HP-ANNs neuronal units exhibit between-attributes spill-over effects. To address this question, we regressed the activity of integration units against prospective gains and losses within temporal windows (i.e. across trials belonging to the same temporal window). The lower-left panel of Figure 5 shows that the sign-rectified sensitivity of HP-ANN units to prospective losses correlates negatively with the range of prospective gains (wide range group: r=-0.06, p<10^-4^; narrow range group: r=-0.05, p<10^-4^). Lastly, we asked whether HP-ANNs reproduce the known range adaptation results of value-sensitive OFC neurons (Padoa-Schioppa, 2009). The lower-right panel of Figure 5 shows that the sign-rectified sensitivity to HP-ANN units to gambles’ expected value correlates negatively with the range of gambles’ expected values (wide range group: r=-0.13, p<10^-4^; narrow range group: r=-0.15, p<10^-4^).

Finally, we ask whether the ANNs reproduce other important features of value-coding neurons in the OFC. It is known that OFC neurons are notoriously diverse in their response profile, but a consistent finding is that, in the context of value-based decision making, they can be classified in terms of so-called “choice cells”, “chosen value cells” and “offer value cells” (Padoa-Schioppa and Assad, 2006, 2008). Given that this can be considered a pre-requisite for any computational model of value integration in the OFC, we asked whether HP-ANNs re produce this known property of OFC neurons.

For each subject, we thus tested whether the response of integration units correlates (across trials) with choice, chosen value and/or gamble value, where value is defined as the weighted sum of gains and losses (according to the static logistic model parameter estimates). Integration units are then classified in terms of which variable it correlates most with (but units are attributed to none of these labels if the correlation is not significant). The results of this analysis are summarized on Figure 6 below.

**Figure 6:**
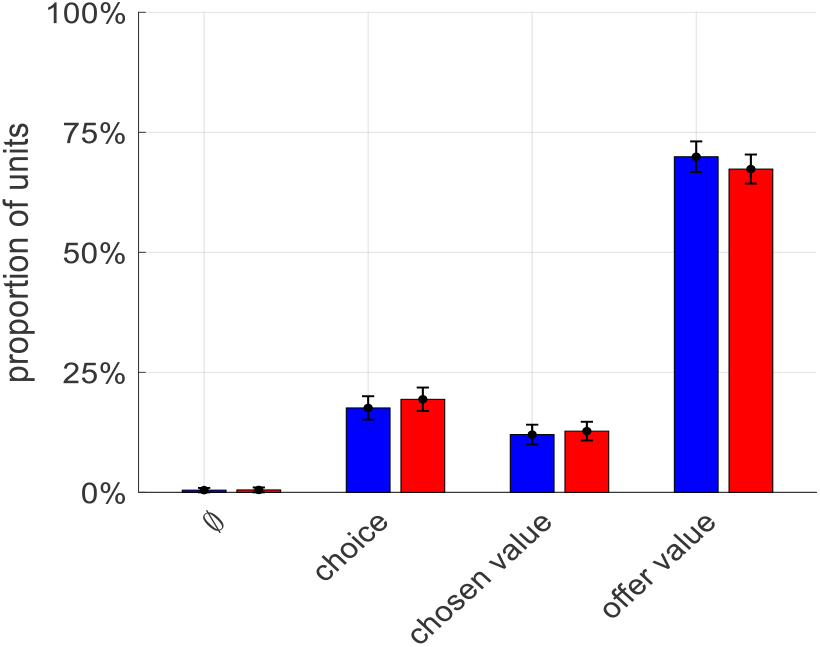
Response profile diversity in HP-ANNs’ integration units. The average proportion of HP-ANNs integration units (y-data) that shows a significant correlation with choice, chosen value, and offer value (or none of these) is plotted for for both groups (blue: wide range, red: narrow range). Errorbars depict s.e.m. (across participants).

It turns out that, although HP-ANN’s integration units were not at all designed to encode these quantities, they eventually reproduce the response variability observed in OFC neurons. This provides additional neurobiological validity to HP-ANN models of range adaptation in the OFC. Importantly, the response profiles are almost identical for both groups. In particular, we find that about 13% of integration units are classified as “chosen value” cells. This is interesting because, under the ANN model, the computational role of these units is in fact exactly the same as that of units that are classified as “offer value” cells: together, they form a population code for the subjective value of gambling. In other words, this classification may not be directly relevant for guessing the underlying computational role of integration units.

## Discussion

In this work, we identify the neural range adaptation mechanism in OFC neurons that mediate the irrational context-dependency of value-based decisions. We focus on decisions under risk, where value needs to be constructed out of primitive attributes (here: prospective gains and losses). This eventually disambiguates the neural and behavioral implications of candidate computational scenarios for range adaptation. We show that Hebbian plasticity between attribute-specific and integration neurons best predicts (out-of-sample) both context-dependent behavioral biases and range adaptation in OFC neurons.

The processing of reward signals in OFC neurons is known to exhibit range adaptation (Conen and Padoa-Schioppa, 2019; Louie and Glimcher, 2012; Rangel and Clithero, 2012). This is typically explained in terms of efficient coding: OFC neurons may adapt their output firing properties to match the recent history of values (Polanía et al., 2019). Implicit in this reasoning is the idea that OFC neurons are receiving value signals, which they are transmitting to downstream decision systems (Padoa-Schioppa and Rustichini, 2014; Rustichini et al., 2017). However, this assumption is at odds with the notion that OFC neurons is rather constructing value from input signals about decision-relevant attributes (O’Doherty et al., 2021; Pessiglione and Daunizeau, 2021b; Raghuraman and Padoa-Schioppa, 2014). In contrast, recent theoretical arguments suggest that flexible, context-dependent, attribute integration in OFC neurons may necessitate Hebbian plastic changes in the synaptic gain of upstream attribute-specific neurons (O’Doherty et al., 2021). Here, we show that these changes eventually translate the distribution of reweighted attribute signals towards the responsive range of integration neurons, incidentally inducing range adaptation. On the behavioral side of things, this scenario predicts between-attribute spill-over effects, such as the impact of contextual gain ranges on peoples’ sensitivity to losses (see Figure 7 in the Methods section). This is important, because such spill-over effects may confound the relationship between range adaptation in OFC neurons and the preference shifts that one might expect under simpler efficient coding scenarios. In this sense, our results complement and extend previous computational modelling studies that focus on the behavioural impact of –spatial-range adaptation in neurons that process specific decision-relevant attributes (Soltani et al., 2012). Interestingly, our results show that, prior to gain range manipulations, people exhibit loss aversion (cf. lower-left panel of Figure 2). Given that loss aversion is –at least partly-determined by temporal range adaptation, this suggests that such stable default attitude may be continuously reinforced by slightly favourable environments. Having said this, many other candidate neural mechanisms may, in principle, compete or interact with Hebbian plasticity, eventually crystalizing or destabilizing plastic changes. For example, homeostatic plasticity is known to induce slow neural hysteretic effects (Fox and Stryker, 2017; Pezzulo et al., 2015; Toyoizumi et al., 2014; Turrigiano, 2017). To what extent these or similar kinds of neurophysiological mechanisms may explain enduring attitudinal and/or behavioral alterations is an open and challenging issue.

**Figure 7:**
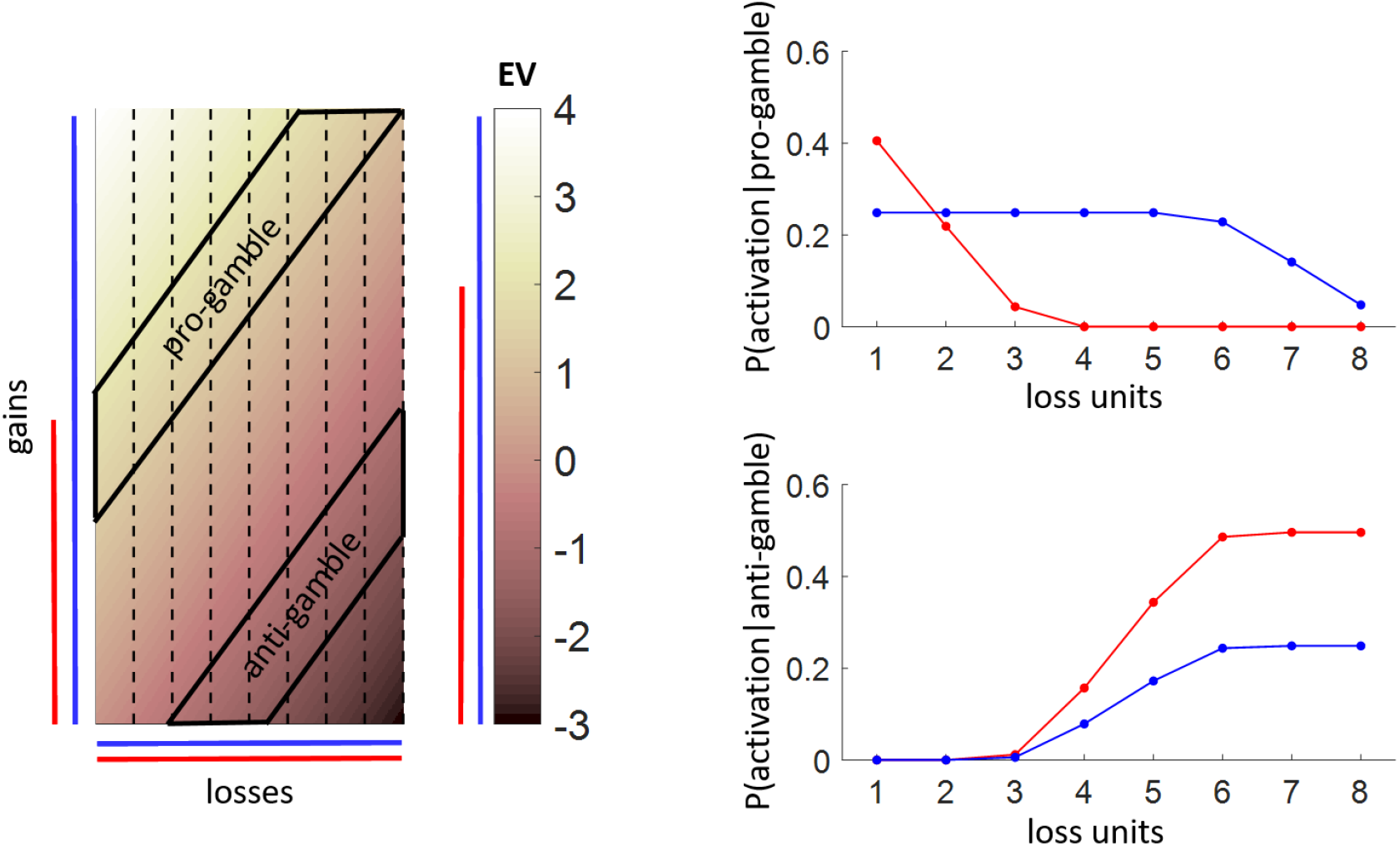
range adaptation under Hebbian plasticity. Left: a schematic representation of the decision value landscape (color code from EV=-3 to EV=4), which is spanned by prospective gains (y-axis) and losses (x-axis). Colored bars depict the range of gains, losses and ensuing gamble value in two different contexts: wide range of gains (blue) and narrow range of gains (red), respectively. Black dotted lines denote the effective frontier of loss domains within which loss-specific units respond to (here: 8 units with pseudo-gaussian activation functions). Black trapezes mark the responsive domains of “anti-gamble” units (which respond when −2<EV<-1) and “pro-gamble” units (which respond when 1<EV<2), respectively. Upper-right: the probability of activation of each loss unit (x-axis) is plotted for each context, when “pro-gamble” units activate. Upper-right: same as upper-right panel, but when “anti-gamble” units activate.

On the neural side of things, it is reassuring to see that fMRI patterns of activity in the lateral OFC strongly resemble the quantitative predictions of HP-ANN models. One might find disappointing that these predictions turn out not to be verified in the vmPFC (at least not in the narrow range group). In fact, there is an ongoing debate regarding the relative contribution of these two OFC subregions w.r.t. value processing. For example, lateral, but not medial, OFC may host representations of attributes that presumably compose value judgements (Suzuki et al., 2017). That we eventually also identify the Striatum and the Amygdala to be involved in this context is well aligned with the existing literature. On the one hand, the ventral Striatum is known to encode value and risk (Schultz et al., 2008), and the tendency to opt for a risky choice increases with the magnitude of the striatal response to risk (Christopoulos et al., 2009; Kuhnen and Knutson, 2005). In fact, the exact same experimental protocol as for the narrow range group already served to demonstrate that the differential striatal responses to losses and gains drive inter-individual variations in loss aversion (Tom et al., 2007). On the other hand, it was also shown that the prospect of a loss prospect may activate Amygdala, which would trigger a cautionary brake on behavior that facilitates loss aversion (Martino et al., 2006, 2010). How OFC, ventral Striatum and Amygdala eventually interact with each other to determine loss aversion is unknown, and the present study does not resolve this debate.

Now, how generalizable is the neural mechanism we disclose here? We argue that synaptic plasticity (and in particular: Hebbian plasticity) may explain many forms of persistent irrational behavioral changes, through temporal range adaptation effects in OFC neurons. We note that, in our context, these changes seem to unfold over several minutes (cf. lower-left panel of Figure 2), which is consistent with the fastest time scale of long-term potentiation/depression (Abraham, 2003). However, we contend that the evidence we provide here is insufficient to establish whether these changes remain stable over longer periods and whether they can be overcome by explicit instructions or intensive training (see, e.g., Cicchini et al., 2012 for an example in the perceptual domain). A related issue is whether similar synaptic plasticity mechanisms may explain virtually instantaneous range adaptation in value-coding neurons (Louie et al., 2015), eventually driving behavioral phenomena such as the framing effect. Here, we speculate that the framing of decisions may automatically trigger contextual expectations regarding expected gain and/or loss ranges, which may induce fast synaptic plasticity within value-constructing networks through, e.g., short-term potentiation (Fiebig and Lansner, 2017).

That the brain’s biology is to blame for all kinds of cognitive and/or behavioural flaws is not a novel idea (Buschman et al., 2011; Marois and Ivanoff, 2005; Miller and Buschman, 2015; Ramsey et al., 2004). However, providing neuroscientific evidence that a hard-wired biological constraint shapes and/or distorts the way the brain processes information is not an easy task. This is because whether the brain deviates from how it *should* process a piece of information is virtually unknown. This is particularly true for value-guided decision making, which relates to subjective assessments of preferences rather than objective processing of decision-relevant evidence (Rangel et al., 2008). Nevertheless, value-guided decision making is known to exhibit many irrational biases, the neurocognitive explanations of which have been the focus of intense research over the past decades. From a methodological standpoint, our main contribution is to show how to leverage computational models (in particular: ANNs) to test hypotheses regarding neurophysiological mechanisms that may constrain or distort behaviorally-relevant information processing. On the one hand, we retain the simplicity of established “model-based” fMRI approaches (Borst et al., 2011; O’Doherty et al., 2007), which proceed by cross-validating the identification of hidden computational determinants of behavior with neural data. On the other hand, our dual ANN/RSA approach enables us to quantify the statistical evidence for neurophysiological mechanisms that are difficult –if not impossible-to include in computational models that are defined at Marr’s *algorithmic* level (McClamrock, 1991), e.g., normative models of behavior (as derived from, e.g., learning or decision theories) and/or cognitive extensions thereof. Hebbian plasticity is a paradigmatic example of what we mean here. Recall that it was initially proposed as an explanation -at the neural or Marr’s *implementational* level- for learning, memory and sensory adaptation (Hebb, 1950). Since then, Hebbian-like synaptic plasticity that serves well-defined computational purposes of this sort has been superseded by theoretical frameworks that transcend the three Marr’s analysis levels, e.g. the “Bayesian brain” hypothesis and related predictive and/or efficient coding scenarios (Aitchison and Lengyel, 2017; Doya et al., 2007; Friston, 2012). But hard-wired biological mechanisms similar to Hebbian plasticity may not always be instrumental to the cognitive process of interest. In turn, it may be challenging to account for incidental biological disturbances of neural information processing, when described at the algorithmic level. A possibility here is to conceive of these disturbances as some form of random noise that perturbs cognitive computations (Drugowitsch et al., 2016; Wyart and Koechlin, 2016). In contrast, we rather suggest relying on computational models that solve a well-defined computational problem (here: constructing the gambles’ subjective value from prospective gains and losses) but operate at the neural level. Accounting for possibly incidental, biological constraints and/or hard-wired mechanisms then enables comparing quantitative/deterministic scenarios for sub-optimal disturbances of covert cognitive processes of interest.

## Methods

### Model-free analysis of behavioral data

First, we describe peoples’ risk attitude in terms of the probability of gambling given the gamble’s expected value EV = 0.5*(G-L), where G and L are the gamble’s prospective gain and loss, respectively. For each participant, we binned trials according to deciles of EV, and measured the rate of gamble acceptance (cf. Figure 2, upper-left panel).

Second, we regressed peoples’ decision to gamble onto gains and losses. Within each participant, we fit the following logistic regression model: p(g_i_) = s(w_0_ + w_G_*G_i_ – w_L_*L_i_), where gi is the binary gamble decision at trial *i,* G_i_ and L_i_ are the prospective gain and loss of trial *i,* w_0_ is the intercept, and w_G_ and w_L_ are the sensitivity to gains and losses, respectively. We then report within-subject parameter estimates at the group-level for random effect analyses (empirical histograms of the ensuing parameter estimates can be eyeballed on Figure S2 of the Supplementary Material). Logistic model parameters can also be recombined to measure peoples’ loss aversion (log(w_L_/w_G_)) and sensitivity to expected value (2*w_G_+2*w_L_). Note that the logistic model can also be used to perform counterfactual model simulations. For each subject, we use the corresponding fitted parameters to evaluate the trial-by-trial probability of gamble acceptance that would have been observed, had this subject/model be exposed to the sequence of prospective gains and losses that each subject *of the other group* was exposed. It turns out that such out-of-sample predictions of peoples’ behaviour are (expectedly) inaccurate. More precisely, such logistic regression cannot predict the observed group-difference in peoples’ risk attitude (see Figure S1 of the Supplementary Material).

Third, we performed a sliding window analysis: decisions were first partitioned into chunks of 16 consecutive trials each, which were then regressed against corresponding gains and losses using the and the same logistic model as above. From this, we obtain a set of logistic parameter estimates (intercept and sensitivity to gains and losses) per temporal window, per subject. Note that the group-averaged temporal dynamics of logistic parameter estimates can be eyeballed on Figure S3 of the Supplementary Material. Temporal changes in the ensuing loss aversion index can thus be followed as time unfolds (cf. lower-left panel of Figure 2). Finally, we measure the effective range of gain and loss attributes per temporal window in terms of their sample variance across trials (within a temporal window). Importantly, given that sequences of prospective gains and losses are randomized across subjects, the temporal changes of attributes ranges is specific to a given participant. We then evaluate the statistical relationship between peoples’ sensitivity to each attribute and attribute ranges, across temporal windows (cf. lower-right panel of Figure 2).

### Artificial neural network models of risk attitude

Artificial Neural Networks or ANNs essentially attempt to decompose a possibly complex form of information processing in terms of a combination of very simple computations performed by connected ‘units’, which are mathematical abstraction of neurons. Here, we take inspiration from a growing number of studies that use ANNs as mechanistic models of neural information processing (Güçlü and Gerven, 2015; Kietzmann et al., 2017, 2019; Kriegeskorte and Golan, 2019), with the added requirement that they have to explain observed behavioural data.

In abstract terms, any decision can be thought of as a cognitive process that transforms some input information *u* = {*u*^(1)^, *u*^(2)^,…*u*^(*n_u_*)^} into a behavioral output response *r*. Here, participants have to accept or reject a risky gamble composed of a 50% chance of obtaining a gain G and a 50% chance of experiencing a loss *L*, i.e. *u* is composed of *n_u_* = 2 input attributes: *u* = {*G,L*}. Under an ANN model of such decisions, people’s behavioral response is the output of a neural network that processes the attributes *u*, i.e.: *r* ≈ *g_ANN_* (*u, ϑ*,) where *ϑ* are unknown ANN parameters and *g_ANN_* (•) is the ANN’s input-output transformation function. So-called “shallow” ANNs effectively reduce *g_ANN_* (•) to a combination of neural units organized in a single hidden layer. In what follows, we rather rely on (moderately) deep ANNs with two hidden layers: namely, an attribute-specific layer (which is itself decomposed into gain-specific and loss-specific layers) and an integration layer (which receives input from both attribute-specific layers). The latter then collectively contribute to gamble decisions by integrating prospective gains and losses.

We assume that each attribute 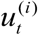 is encoded into the activity of neurons 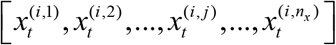 of its dedicated “attribute-specific layer”, where *n_x_* is the number of aatribute-specific neurons per attribute. What we mean here is that the neuron *j* in the attribute-specific layer *i* responds to 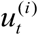 as follows:

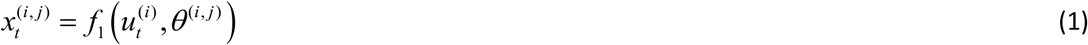

where *f*_1_(•) is the activation function of neural units that compose the ANN’s attribute-specific layer. Collectively, the activity vector 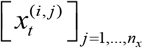 forms a multivariate representation of attribute 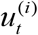 in the form of a population code (Ebitz and Hayden, 2021).

Critically, we consider activation functions that are bounded, i.e. units’ outputs follow either a sigmoidal or a pseudo-gaussian mapping of inputs (see below):

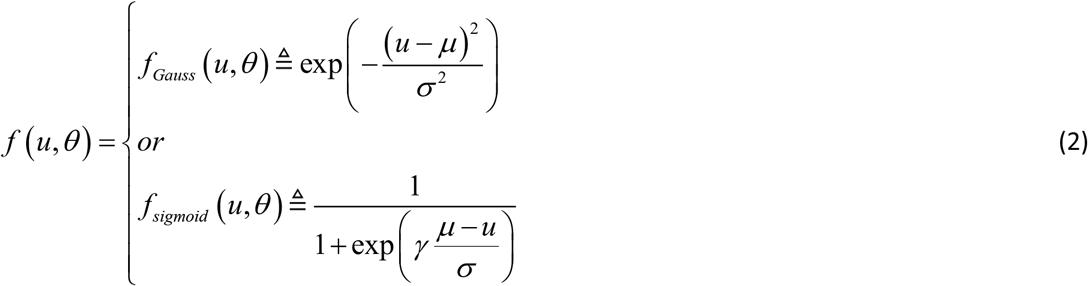

where *γ* ≈ 1.5434 is a scaling constant that we introduce for mathematical convenience (see Supplementary Material). The parameters *θ*^(*i,j*)^ = {*μ*^(*i,j*)^, *σ*^(*i,j*)^} capture idiosyncratic properties of the neuron *j* in the input layer *i* (e.g., its firing rate threshold *μ*^(*i,j*)^ and the pseudo-variance parameter *σ*^(*i,j*)^). Note that, when inputs *u* fall too far away from *μ* (say outside a 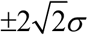 range), both these activation functions saturate, i.e. they produce non discriminable outputs (close to 0 or 1). In other words, the pseudo-variance parameter defines the range of inputs over which units incur no information loss.

Then the output of the attribute-integration layers is passed to the “integration layer” 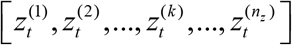, i.e. the neuron *k* of the integration layer responds to 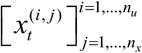 as follows:

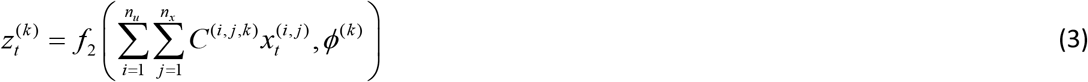

where *C*^(*i,j*)^ is the connection weight from the neuron *j* in the attribute-specific layer *i* to the neuron *k* of the integration layer, and *ϕ*^(*k*)^ capture idiosyncratic properties of the integration neuron *k*. Without loss of generality, we set *n_z_* = *n_x_*.

The behavioral response *r_t_* at time or trial *t* is then read out from the integration layer as follows:

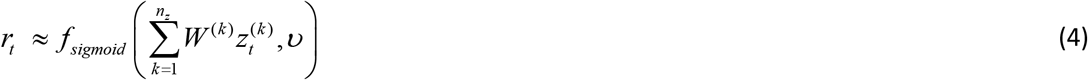

Collectively, integration neurons form a representation of decision value in the form of a population code of decision value, which is read through the weights’ pattern *W*^(*k*)^. Taken together, Equations 1-3-4 define the end-to-end ANN’s transformation of prospective gains and losses into gamble decisions:

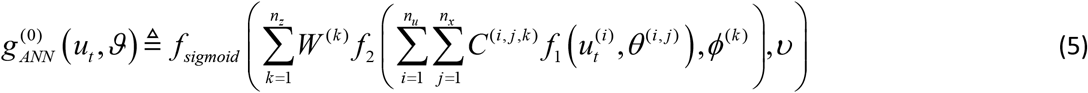

where *ϑ* lumps all ANN parameters together, ie.: 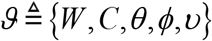, and *f*_*i*∈{1,2}_ are either gaussian or sigmoid. A schematic summary of the ANN’s double-layer structure is shown on the upper-left panel of Figure 3.

Here, we refer to this default ANN architecture as ‘static’ ANN. Note that, provided there are enough neurons in attribute-specific and integration layers, this ANN architecture can capture any value function defined on the bidimensional input space spanned by prospective gain/loss pairs. However, it cannot capture range adaptation effects, whereby the recent history of prospective gains and losses change the value population code. This is why we now introduce hebbian plasticity.

The ANN’s structure enables the explicit modelling of Hebbian plasticity between attribute-specific and integration layers. More precisely, the Hebbian adaption rule will strengthen the connection between attribute-specific and integration units that covary across decision trials. This recapitulates the “fire together, wire together” rule:

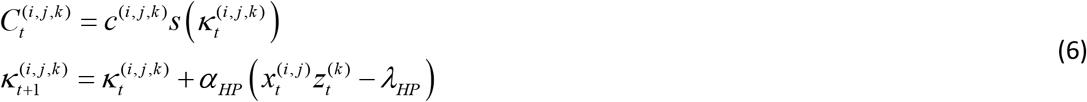

where *c*^(*i,j,k*)^ and 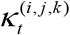 are the static and dynamic components of between-layers connection weights, respectively, *α_HP_* is the Hebbian update rate and *λ_HP_* is the Hebbian covariance threshold. Equation 6 reinforces a connection weight whenever the product of the corresponding units’ outputs exceeds the threshold *λ_HP_*. We coin ANNs modified with Equation 6 ‘HP-ANN’.

In our context, Hebbian plasticity between attribute-specific and integration units is motivated by the necessity, for OFC integration neurons, to flexibly select context-dependent attributes (O’Doherty et al., 2021). However, by translating the reweighted outputs of attribute-specific units within the responsive range of integration units, its incidental effect is to induce some form of temporal range adaptation. In particular, it will induce between-attribute spill-over effects (see Figure 7 below).

For the sake of simplicity, we focus on pseudo-gaussian activation functions, here. This implies that loss-specific units will respond to a specific domain of prospective losses. Similarly, integration units would respond to a specific domain of gamble values. Consider integration units that respond when 1<EV<2. In both contexts, they can be thought of as “pro-gamble” units, in that they signal a rather high gamble value. In the wide range context, they will covary with almost all loss units, whereas in the narrow range context, they will unlikely covary with loss units that signal high losses. In turn, Hebbian plasticity will render the connection between “pro-gamble” units and “high loss” units stronger in the wide range context than in the narrow range context. In other terms, those integration units that signal high gamble values become more sensitive to losses in the wide range context than in the narrow range context. The effect is reversed for “anti-gamble” units (integration units that respond when - 2<EV<-1), which become less sensitive to losses in the wide range context than in the narrow range context. Taken together, Hebbian plasticity will thus tend to desensitize gamble value from losses in the wide range context, when compared to the narrow range context.

At the limit when update rates tend to zero (*α_HP_* → 0), HP-ANNs exhibit no range adaptation, i.e. they become indistinguishable from the above ‘static’ ANNs. Otherwise, hebbian plasticity makes the ANN’s trial-by-trial response a function of the recent history of inputs to the network. Here, update rates effectively control the time scale at which range adaptation will operate. Given that the sequence of prospective gains and losses is arbitrary, different update rates will eventually induce different distortions of trial-by-trial behavioral and/or neural responses. Therefore, to capture idiosyncrasies in behavioural range adaptation, we need to estimate update rates (along with other ANN parameters) from peoples’ observed trial-by-trial gamble responses. Here, we rely on established variational Bayesian model inversion techniques to perform probabilistic parameter estimation (Daunizeau, 2017; Friston et al., 2007). To mitigate the impact of local optima, we use a twofold strategy. First, we concurrently fit the behavior trial series together with its rolling mean and variance (over a sliding temporal window whose width we set to 5 trials). Second, we use a hierarchical group-level mixed-effects approach that constrains subject-specific parameter estimates with empirical group statistics (Daunizeau, 2019). The priors on the ANNs’ model parameters for the ensuing parametric ‘empirical Bayes’ approach are summarized in Table S1 of the Supplementary Material. In addition, empirical histograms of HP-ANN’s threshold and update rate parameters can be eyeballed on Figure S5. We performed all the data analyses using the VBA academic freeware (Daunizeau et al., 2014) : https://mbb-team.github.io/VBA-toolbox/.

For each subject and each ANN model (6 models: 3 ANN structures and 2 kinds of activation functions), this procedure provides (i) ANN parameter estimates, (ii) trial-by-trial postdictions of gamble acceptance, and (iii) trial-by-trial predictions of neural patterns within both attribute-specific and integration layers. From a statistical perspective, Equations 6 and 7 provide extra degrees of freedom when fitting HP-ANNs to behavioural choices, when compared to static ANNs. This means that one would expect fit accuracy to be higher for ANNs with hebbian plasticity, irrespective of whether this mechanism is a realistic determinants of behavior or not (see Figure S6 in the Supplementary Material). This is why we cross-validate all our behavioral analyses. Importantly, each set of within-subject fitted parameters can be used to derive counterfactual predictions of the series of decisions that would have been obtained, had the corresponding participant been exposed to another sequence of prospective gains and losses. For each participant, we thus derived all series of out-of-sample behavioural predictions that correspond to the sequences of gain/loss pairs experienced by participants in the other group. Trials were binned according to EV deciles prior to averaging (twice) over subjects the predicted rate of gamble acceptance, for each model. Out-of-sample behavioural predictions for all ANN models can be eyeballed on Figure 3.

Finally, the postdicted behavioral responses of candidate ANNs can be analysed similarly to observed participants’ decisions (for posterior predictive checks). More precisely, we perform the same logistic regressions, this time on binarized ANNs’ decision outputs (against prospective gains and losses). This enables us to derive ANN-based within-subject loss aversion indices, both globally and across sliding temporal windows. In addition, we assess the statistical relationship between integration units’ sensitivity to gains and losses and gain/loss ranges, across temporal windows (for each candidate ANN). To reproduce typical electrophysiological analyses (Padoa-Schioppa, 2009), we focus on the units’ absolute gain/loss sensitivity to (cf. “sign-rectified” sensitivities). The results of these analyses for HP-ANNs are shown on Figure 5 of the main text.

### fMRI data: experimental paradigm, pre-processing and univariate analyses

In this work, we perform a re-analysis of the NARPS dataset (Botvinik-Nezer et al., 2019, 2020), openly available on *OpenNeuro.org* (Poldrack et al., 2013). This dataset includes two studies, each of which is composed of a group of 54 participants who make a series of decisions made of 256 risky gambles. On each trial, a gamble was presented, entailing a 50/50 chance of gaining an amount G of money or losing an amount L. As in Tom et al. (2007), participants were asked to evaluate whether or not they would like to play each of the gambles presented to them (strongly accept, weakly accept, weakly reject or strongly reject). They were told that, at the end of the experiment, four trials would be selected at random: for those trials in which they had accepted the corresponding gamble, the outcome would be decided with a coin toss and for the other ones -if any-, the gamble would not be played. In the first study (hereafter: “narrow range” group), participants decided on gambles made of gain and loss levels that were sampled from the same range (G and L varied between 5 and 20 $). In the second study (hereafter: the “wide range” group), gain levels scaled to double the loss levels (L varied between 5 and 20$, and G varied between 10 and 40$). In both studies, all 256 possible combinations of gains and losses were presented across trials, which were separated by 7 seconds on average (min 6, max 10).

MRI scanning was performed on a 3T Siemens Prisma scanner. High-resolution T1w structural images were acquired using a magnetization prepared rapid gradient echo (MPRAGE) pulse sequence with the following parameters: TR = 2530 ms, TE = 2.99 ms, FA = 7, FOV = 224 × 224 mm, resolution = 1 × 1 × 1 mm. Whole-brain fMRI data were acquired using echo-planar imaging with multi-band acceleration factor of 4 and parallel imaging factor (iPAT) of 2, TR = 1000 ms, TE = 30 ms, flip angle = 68 degrees, in plane resolution of 2X2 mm 30 degrees of the anterior commissure-posterior commissure line to reduce the frontal signal dropout, with a slice thickness of 2 mm and a gap of 0.4 mm between slices to cover the entire brain. See https://www.narps.info/analysis.html#protocol for more details. Data were preprocessed using SPM (https://www.fil.ion.ucl.ac.uk/spm/), following standard realignment and movement correction guidelines. Note that we excluded 5 participants from the narrow range group because the misalignment between functional and anatomical scans could not be corrected.

We conducted two sets of univariate analyses, using two distinct general linear models or GLMs (Friston et al., 1995). All models incorporated only one event per trial, which was the onset of gamble presentation. In the first GLM, we used two parametric modulations of the trial epoch regressor: namely prospective gains (G) and losses (L). In the second GLM, we used only one parametric modulation, i.e. the gamble’s expected value EV=0.5*(G-L). To account for hemodynamic delays, all regressors of interest were convolved with a canonical hemodynamic response function, as well as with its temporal derivative (Hopfinger et al., 2000). Potential motion artifacts were explained away by introducing subject-specific realignment parameters which were modeled as covariates of no interest. Linear contrasts of regression coefficients were computed at the subject level, smoothed with an 8mm FWHM kernel, and then taken to a group level random effect analysis using a one-sample t-test. Note that gain, loss and gamble’s expected value GLM regressors where mean-centered but not rescaled (e.g., z-scored), to allow for a proper between-group comparison of neural sensitivity to these factors. Finally, we also performed a conjunction analysis (Friston et al., 2005) to find brain regions that both significantly responded to contrasts of interests (when pooling both groups together) and showed a significant group difference.

### ANN-based representational similarity analysis (RSA)

In the results Section, we focus on three subregions of the OFC: namely, left/right lateral OFC (BA13) and medial OFC (vmPFC). However, our and previous mass-univariate analyses of the same dataset provided evidence for the implication of multiple brain systems in processing prospective gains and losses, in particular: the lateral and medial OFC, the dorsolateral prefrontal cortex or dlPFC, the anterior cingulate cortex or ACC, the posterior cingulate cortex or PCC, the Amygdala, the Striatum and the Insula (Botvinik-Nezer et al., 2020). For the sake of completeness, we thus included all these ROIs in our analyses. We first extracted binarized masks fo each ROI using the NeuroQuery website (Dockès et al., 2020). We then took the 2000-th strongest voxels, excluded those that belonged to clusters smaller than 200 voxels, smooth the resulting map, filter out white matter overlaps, and kept the 200 strongest voxels of each remaining clusters. This procedure yielded approximately spherical ROIs with similar sizes across all ROIs (see Figure 4 and S9). In each ROI, we regressed trial-by-trial activations with SPM through a GLM that included one stick regressor for each trial (at the time of the gamble presentation onset), which was convolved with the canonical HRF. To account for variations in hemodynamic delays, we added the basis function set induced by the HRF temporal derivative. To correct for movement artifacts, we also included the six head movement regressors and their squared values as covariates of no interest. We then extracted the 256 trial-wise regression coefficients in each voxel of each ROI. This effectively provided BOLD trial series *Y_fMRI_* that are deconvolved from the hemodynamic response function (Dale, 1999). No spatial smoothing was applied to preserve information buried in spatial fMRI activity patterns.

Once fitted to behavioral data, each ANN models makes specific trial-by-trial predictions of neural activity patterns {*x_t_,z_t_*} that can be compared to multivariate fMRI signals in each ROI. This enables us to evaluate the biological realism of candidate models. Here, we have chosen to rely on a modified representational similarity analysis or RSA (Diedrichsen and Kriegeskorte, 2017; Friston et al., 2019; Kriegeskorte, 2008). In brief, RSA consists of evaluating the statistical resemblance between model-based and data-based ‘representational dissimilarity matrices’ or RDMs, which we derive as follows. Let *Y* be the *n_y_* × *n_t_*, multivariate time series of (modelled or empirical) neural activity, where *n_y_* and *n_t_* are the number of units and trials, respectively. Note that, for model predictions, ‘units’ mean artificial elementary units in ANNs, whereas for fMRI data, ‘units’ mean voxels in a given ROI. First, we orthogonalize *Y* with respect to potential confounding sources of between-trial variability, i.e.: *Y* ← *Y* (*I_n_t__* – *X^T^* (*XX^T^*)^-1^ *X*), where *X* is the *n_c_* × *n_t_* confounds matrix. Here, the set of confounds typically include a constant term, gains, losses and participants’ gamble decision. Second, we standardize neural time series by zscoring over trials. Third, we derive the *n_t_* × *n_t_* between-trials Euclidean distance matrix *D_Y_* :

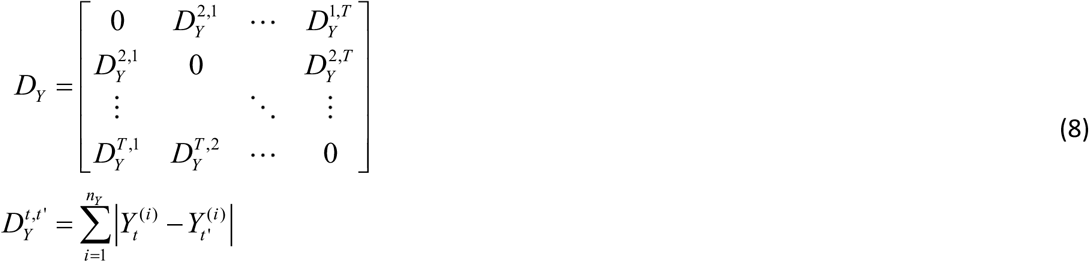

where the matrix element 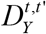 thus measures the dissimilarity of neural patterns of activity between trial *t* and trial *t*′, having removed trial-by-trial variations that can be explained as linear combinations of gains, losses and gamble decisions. Fourth, we proceed with the statistical comparison of *D_Y_ANN__* and *D_Y_fMRI__*, where *D_Y_ANN__* is derived from activity patterns of the ANNs’ integration layer and *D_Y_fMRI__* is *‘fMRI ‘AW ‘fMRI* derived from HRF-deconvolved multi-voxel fMRI trial series in each ROI. We define RDMs as the lower-left triangular part of *D_Y_*. In line with recent methodological developments of RSA (Diedrichsen and Kriegeskorte, 2017), we first bin RDMs into 20 quantiles, and then compute the Pearson correlation *ρ* = corr (*RDM_ANN_, RDM_fMRI_*) between the binned RDMs. We then assess group-level statistical significance of RDMs’ correlations using one-sample t-tests on the group mean of Fischer-transformed RDM correlation coefficients *ρ* (we use the Bonferroni correction for multiple comparisons across ROIs, for each model).

Note that ANN-RSA summary statistics (such as RDM correlation coefficients) *do not* favor more complex ANNs (i.e. ANNs with more parameters, such as HP-ANNs). This is because, once fitted to behavioural data, ANNs produce activity patterns that have no degree of freedom whatsoever when they enter RDM derivations. In particular, this means that static ANNs can a priori show a greater RDM correlation than HP-ANNs. In turn, this enables a simple yet unbiased statistical procedure for comparing candidate models, based on group-level comparisons of RDM correlation coefficients. In particular, we can assess the statistical significance of the comparison of RDM correlation of each candidate ANN against all the other ones. One can show that this procedure is immune to arbitrary modelling choices such as the total number of units in ANN models. Details regarding this procedure are provided in the Supplementary Material. Also, our modified RSA compares multivariate information in ANNs and fMRI data that is orthogonal to linear combinations of decision attributes and behavioral responses. This ensures that the ensuing inferences are orthogonal to previous mass-univariate analyses (cf. selected ROIs respond to gains, losses and/or gambles), and prevents statistical biases towards models that best explain behavioral data.

## Supplementary Material

### 1. Logistic regression of behavioral data: postdiction and out-of-sample predictions

The logistic regression analysis of peoples’ sensitivity to gains and losses provides a trial-by-trial prediction of gamble acceptance, for each pair of prospective gain and loss. But are hard decisions (e.g. where the gamble’s expected value is close to zero) as well explained as difficult decisions (where there is a strong incentive to gamble)? To address this question, we binned trials according to deciles of gambles expected value (see Methods), and measured the rate of the logistic model’s postdiction error (see Figure S1, left panel).

Note: hereafter, we refer to “postdictions” as model predictions on data that was used to fit the model’s parameters. In contrast, “out-of-sample predictions” are model proper predictions on yet unseen data.

**Figure S1:**
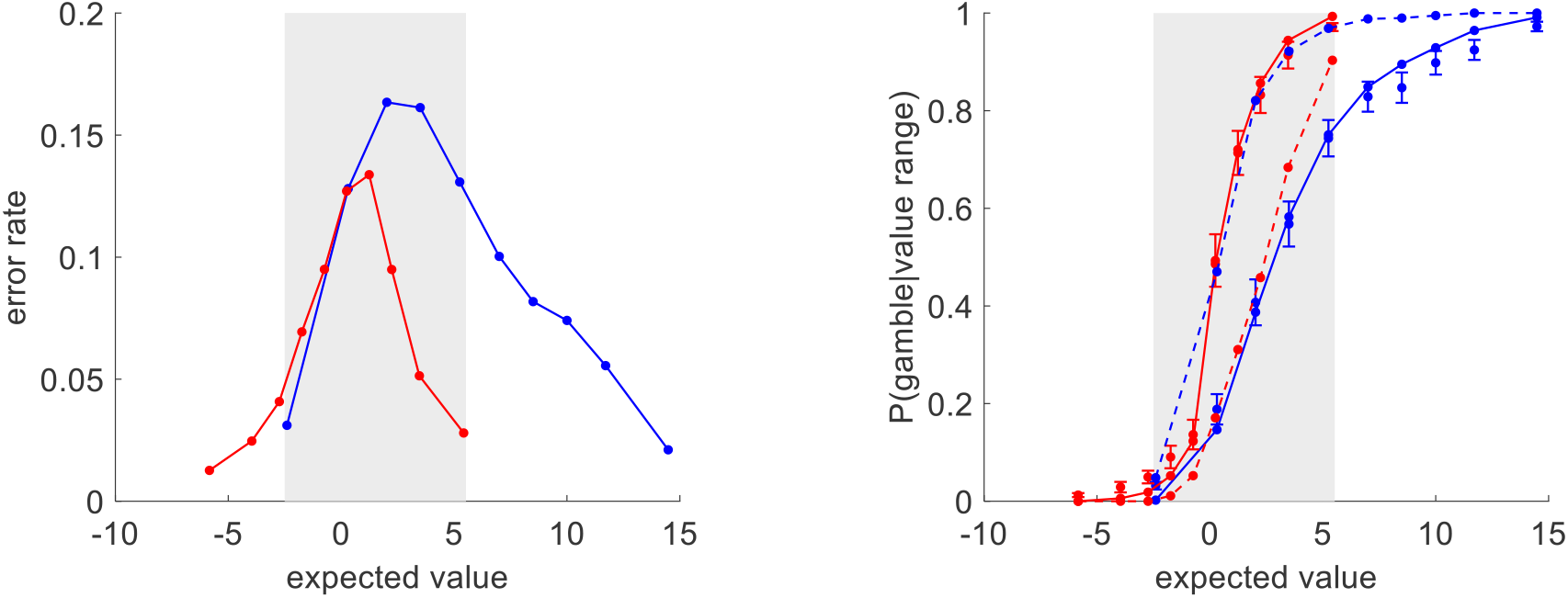
postdiction error and out-of-sample predictions of the logistic model. Left panel: “postdiction” error (y-axis) is plotted against deciles of gambles’ expected value (x-axis), in both groups (red: narrow range, blue: wide range). The grey shaded area highlights the range of expected values that is common to both groups. Right panel: The probability of gamble acceptance (y-axis) is plotted against gambles’ expected value (x-axis), in both groups (red: narrow range, blue: wide range). Dots show raw data (errorbars depict s.em.). Plain and dashed lines show postdictions and out-of-sample predictions, respectively.

One can see that the logistic regression model achieves relatively similar postdiction error profiles in both groups. In particular, easy decisions (either low or high expected value) exhibit much lower inconsistency than difficult decisions (expected value around zero). We also performed counterfactual model simulations: for each subject we simulated the trial-by-trial gamble acceptances that would have been observed, under the logistic model, had this subject/model be exposed to the sequence of prospective gains and losses that each subject of the other group was exposed (see Figure S1, right panel). Those out-of-sample predictions are inaccurate: within the range of expected values that is common to both groups, it wrongly predicts that risk aversion should be lower in the wide range group than in the narrow range group. This is not surprising, because predictions of the logistic model are entirely determined by the pair of prospective gain and loss. In other words, for a given gamble (defined in terms of its prospective gain and loss), the logistic model makes an out-of-sample prediction that corresponds to its equivalent postdiction (or is an extrapolation of it, if performed outside the range of expected value it was originally trained with).

### 2. Logistic regression of behavioral data: empirical distributions of parameter estimates

In the main text, we provide summary statistics for the group differences in the logistic regression parameter estimates. Figure S2 below shows the corresponding empirical histograms.

**Figure S2:**
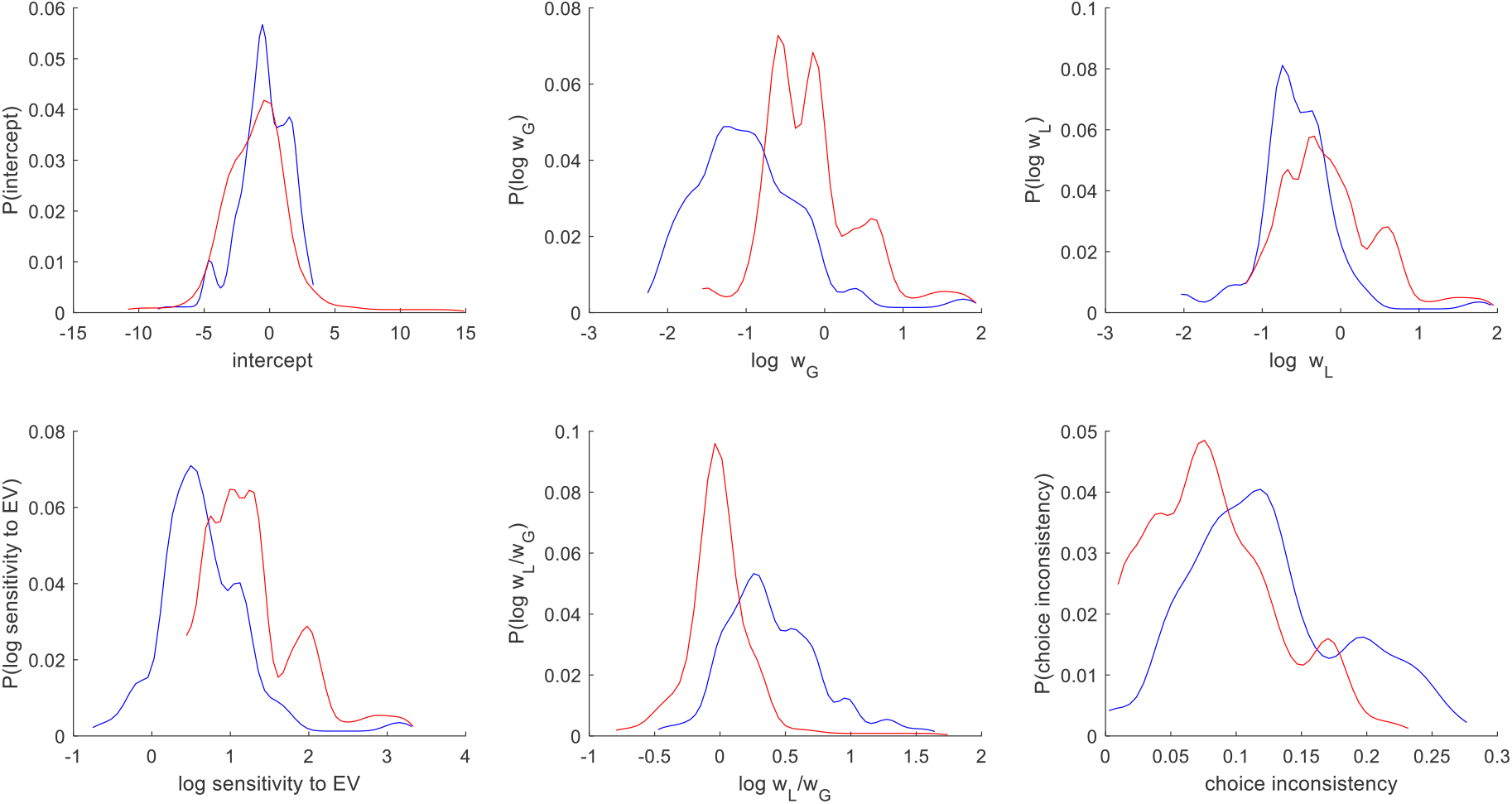
empirical histograms of logistic regression parameters. Upper-left panel: empirical histogram of the intercept parameter for the narrow range group (red) and the wide range group (blue). Upper-middle panel: log-sensitivity to gains (log w_G_). Upper-right panel: log-sensitivity to losses (log w_G_). Lower-left panel: loss aversion index (log w_L_/w_G_). Lower-middle panel: log sensitivity to expected value. Lower-right panel: choice inconsistency.

Upper panels recapitulate Figure 1 of the main text: no group difference w.r.t. intercept, and higher loss and gain sensitivity for people in the narrow range group when compared to people from the wide range group. The lower-left panel shows the ensuing sensitivity to expected value (EV), which is significantly higher for people from the narrow range group than for people from the wide range group (p<10-4, F=20.6, df=101). This is a quantitative summary of the upper-left panel of Figure 1, which depicts the slope of gambling probability w.r.t. to EV for both groups. One can also appreciate inter-individual differences in loss aversion (lower-middle panel of Figure S2). In particular, within the narrow range group, the dispersion of loss aversion is strikingly symmetrical around zero. In contrast, there are very few individuals that show negative loss aversion in the wide range group.

Finally, recall that the rate of choices that deviate from this logistic model provides a simple proxy for individual choice consistency. One can see (cf. lower-right panel of Figure S2) that choice inconsistency is significantly higher in the narrow range group than in the wide range group (p=0.002, t=3.11, df=101). This is important, because choice inconsistency can be (at least partially) explained in terms of a side effect of range adaptation mechanisms. In brief, as neural responses to gain and loss prospects change (as a function of the history of gains and losses), people’s behaviour deviates from any form of static model, including the logistic regression model.

### 3. Logistic regression of behavioral data: sliding window analysis

In the main text, we show the temporal dynamics of loss aversion (cf. Figure 1, lower-left panel). Figure S3 below shows the temporal dynamics of logistic regression parameters, and of the corresponding instantaneous choice inconsistency (i.e. the rate of choice that deviate from the logistic model within each temporal window). One can see that groups mostly differ in terms of sensitivity to gains and losses. More precisely, although both choice inconsistency and the intercept do change over time, they show no group difference (no statistical between-group comparison survives the Bonferroni correction across temporal windows here). In particular, the former result rules out the possibility that the observed between-group difference in risk attitude (cf. Figure 1, upper-left panel in the main text) is due to a differential change in choice stochasticity (behavioural temperature). Note that one trivial possibility could have been that subjects somehow aim at balancing their yes/no decisions, e.g., by gradually shifting their EV threshold to reach about 50% gambles. However, the latter result rules essentially out this possible confound.

**Figure S3:**
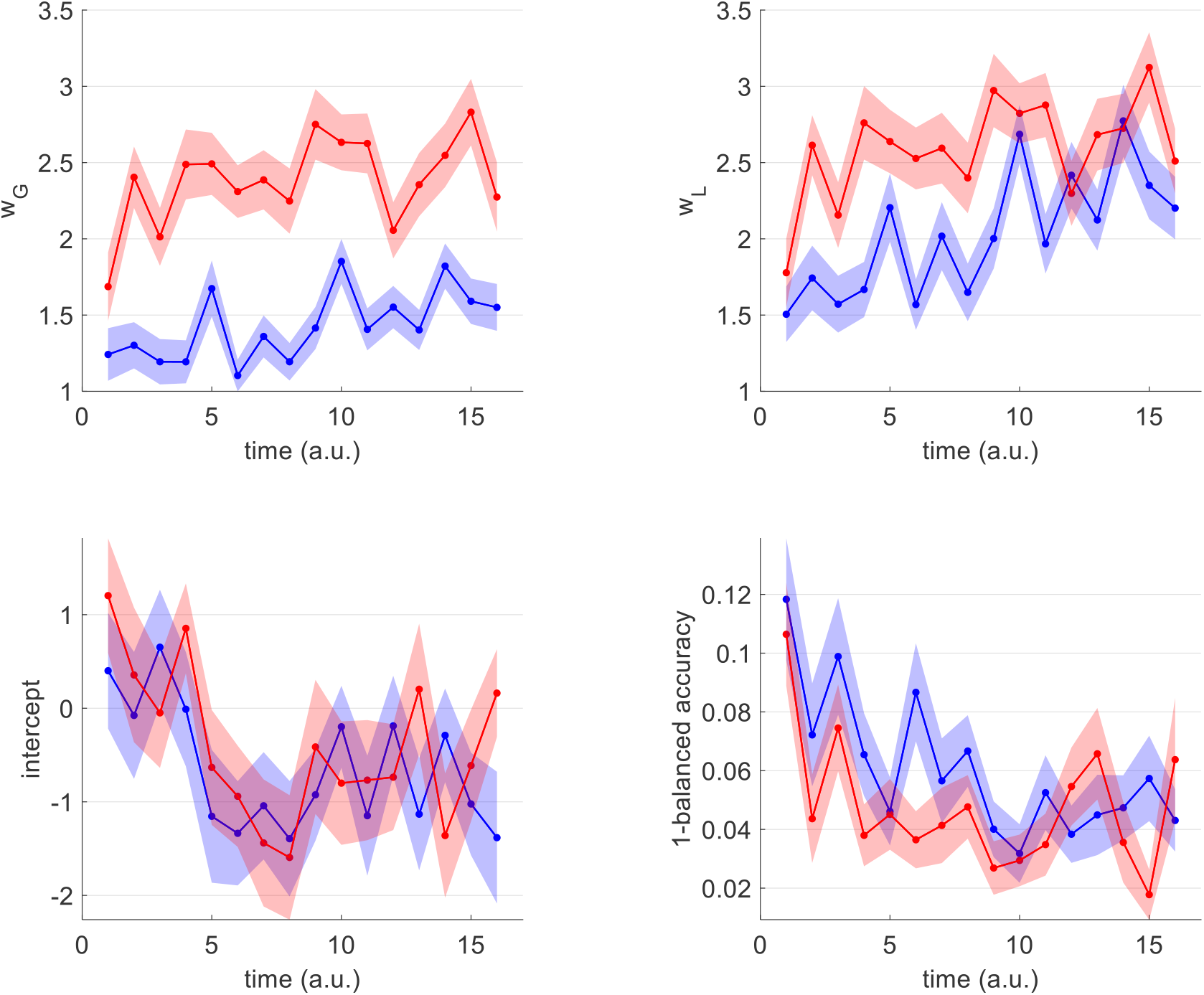
temporal dynamics of logistic regression parameters. Upper-left panel: sensitivity to gains (w_G_, y-axis) is plotted sa a function of time (x-axis) for both groups (red: narrow range, blue: wide range). Plain lines show the group means ans shaded areas depict the s.e.m. Upper-right panel: same as upper-left panel, sensitivity to losses. Lower-left panel: same as upper-left panel, intercept. Lower-right panel: same as upper-left panel, choice inconsistency (measured in terms of 1-balanced accuracy).

Let us now focus on the dynamics of peoples’ sensitivity to gains and losses. In brief, both gain and loss sensitivity increase over time, in both groups. This can be explained by a progressive dampening of apparent choice stochasticity, which would mean that gains and losses would have higher explanatory power, hence the increase in the magnitude of parameter estimates over time. But what is the nature of apparent choice stochasticity? A possibility is that it is due to covert changes in the neural processes that integrate gains and losses to generate the behaviour. Under this assumption, people who show stronger temporal variations in relative gain/loss sensitivities would also exhibit higher apparent choice stochasticity. For each participant, we thus measured the amount of temporal changes in loss aversion in terms of its sample variance across temporal windows. In both groups, choice inconsistency (measured using the static logistic regression model) significantly correlates with changes in loss aversion (narrow range: r=0.41, p=0.003, wide range: r=0.48, p<10^-4^), and this correlation shows no group difference (p=0.40, t=0.84, df=99). If this can be generalized to apparent choice stochasticity within temporal windows, the upper panels of Figure S2 essentially mean that people progressively converge to a static neural processing of behaviorally-relevant information.

### 4. Efficient coding: validity of behavioral predictions

In the main text, we summarize the statistical relationships that exist between temporal changes in peoples’ sensitivity to gains and/or losses and temporal changes in gain/loss ranges (cf. lower-right panel of Figure 1). For the sake of completeness, Figure S4 below shows the 4 sets of statistical relationships, for each group of participants.

**Figure S4:**
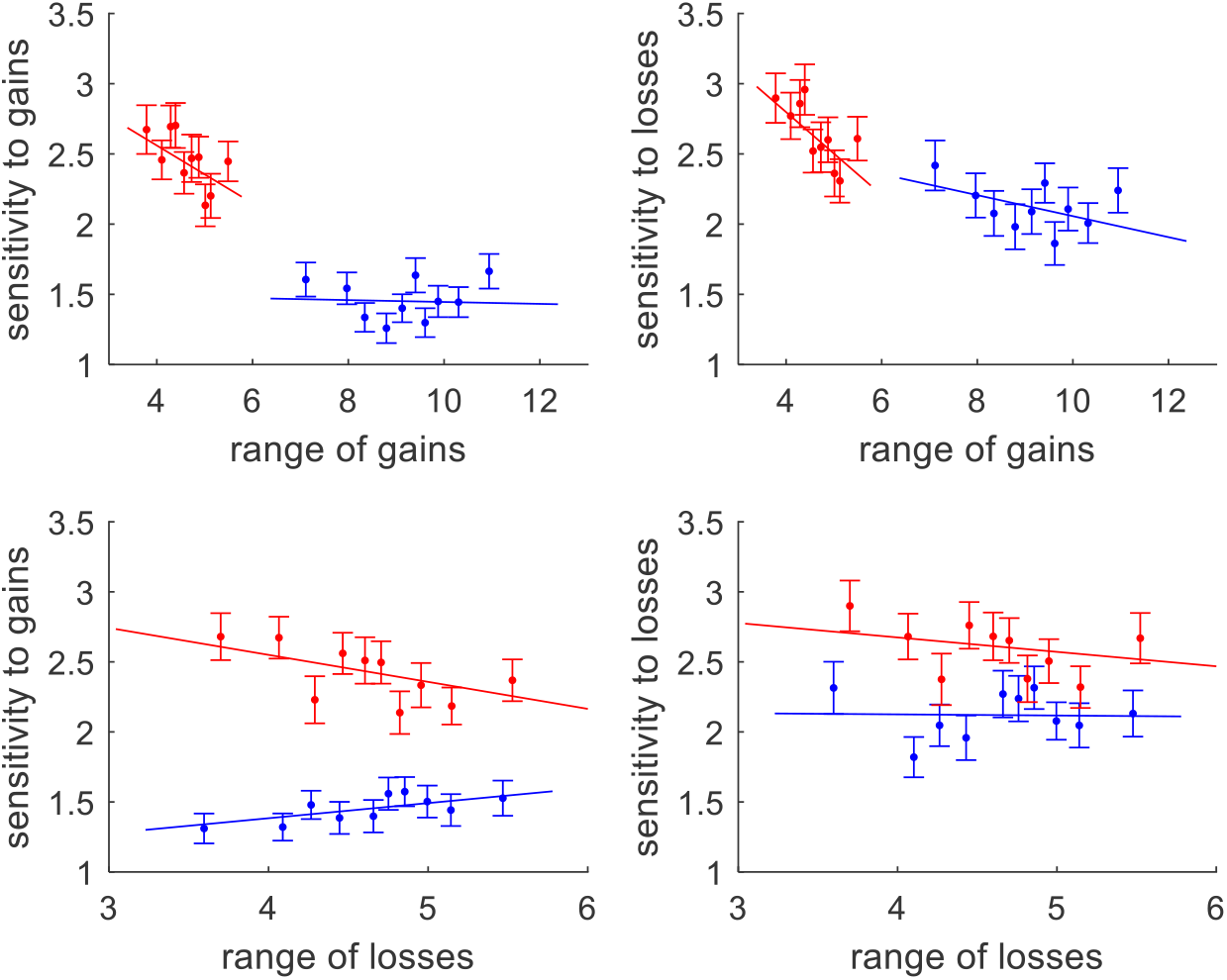
statistical relationships between peoples’ sensitivity to gain/loss prospects and gain/loss ranges. Upper-left panel: sensitivity to gains (y-axis) is plotted against deciles of gain ranges (x-axis, blue: wide range group, red: narrow range group). Upper-right panel: sensitivity to losses against gain ranges. Lower-left panel: sensitivity to gains against loss ranges. Lower-right panel: sensitivity to losses against loss ranges.

### 5. Bayesian priors on ANNs’ parameters

We fit each candidate ANNs to observed trial-by-trial gamble decision sequences using a dedicated Bayesian approach, which requires setting specific prior distributions on model parameters. These priors distributions are summarized on Table S1 below.

**Table S1:**
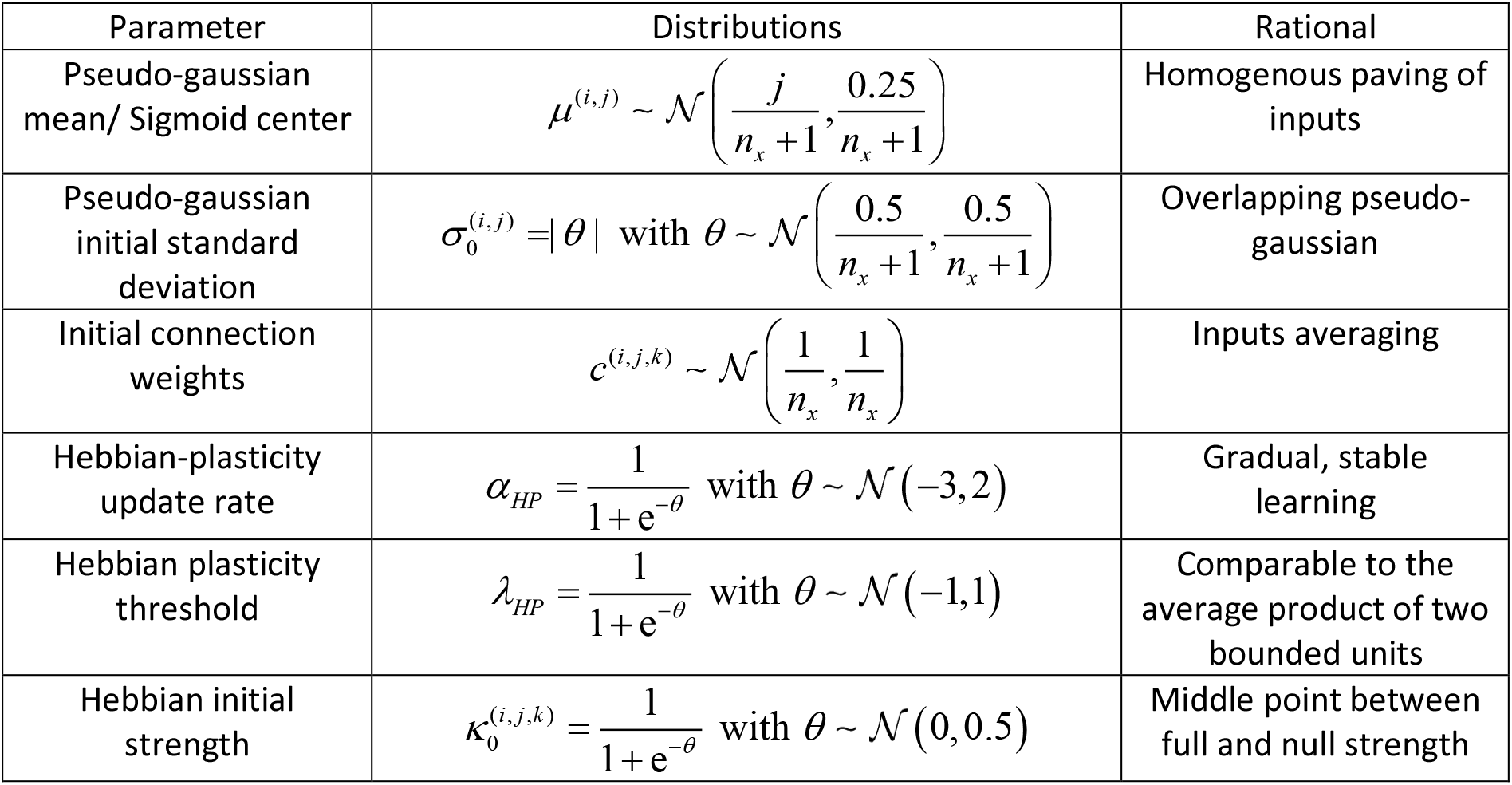
Parameters’ priors for biologically-constrained ANNs.

All parameter notations are defined in the Methods section of the main text. Note that we performed all the analyses using the VBA academic freeware (Daunizeau et al., 2014). Although this toolbox only handles Gaussian prior distributions, native (Gaussian) VBA parameters can be passed through arbitrary mappings prior to entering model computations. This enables VBA to enforce any required constraint (see e.g., Daunizeau, 2017). This is the case here for all model parameters except the centers of units’ activation functions (e.g., HP-ANN update rates need to be constrained between 0 and 1). In Table S1, θ denotes VBA native parameters: they are given Gaussian prior distributions, and then passed through the appropriate nonlinear mapping.

### 6. Parameter estimates of plastic ANNs

Of particular interest in our ANN-based analysis of behavioral data are the parameters that control the update rules of efficient coding and Hebbian plasticity (HP-ANN’s update rates *α_HP_* and covariance thresholds *λ_HP_*). Figure S5 below shows the empirical histograms of parameters estimates, for both groups of participants.

**Figure S5:**
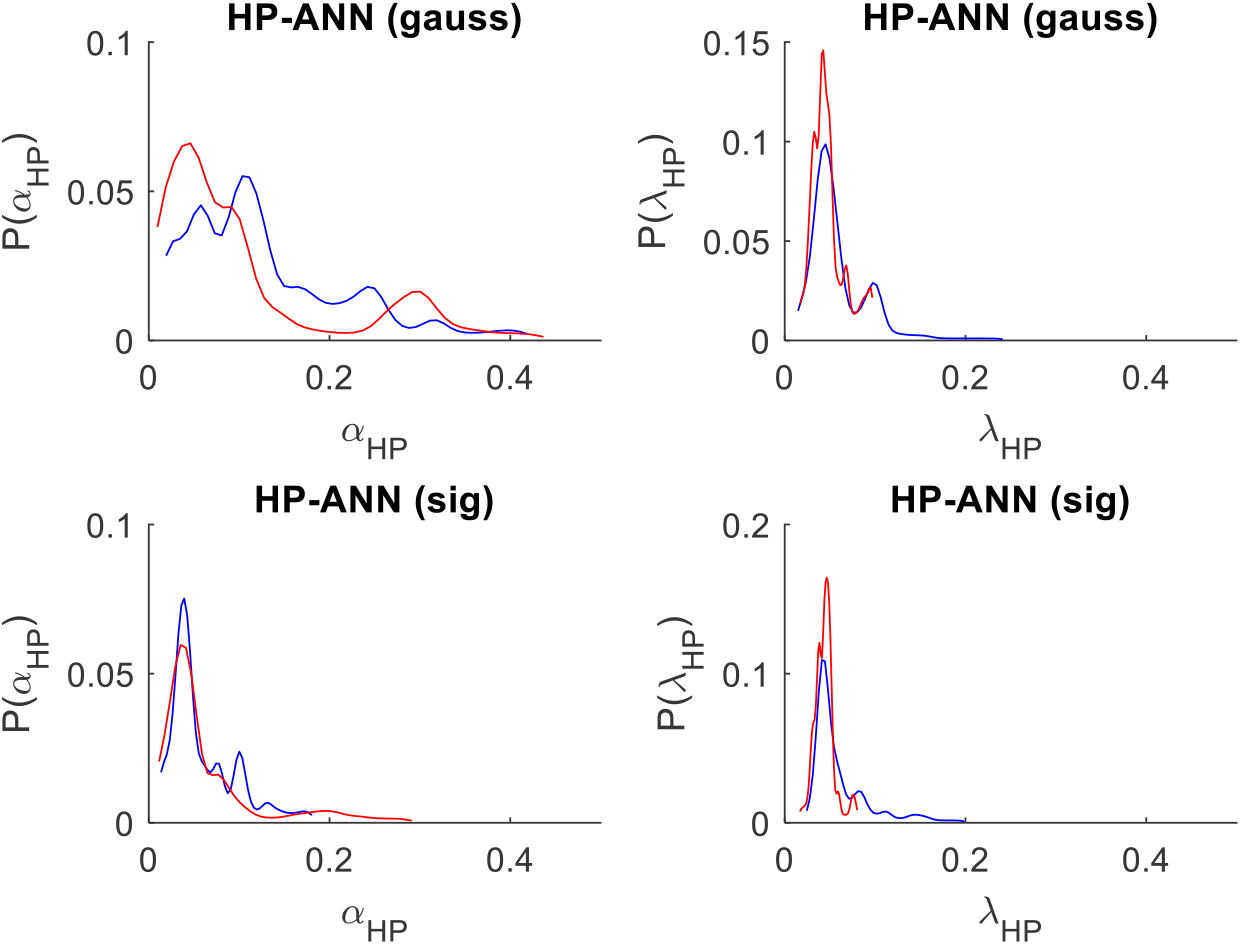
HP-ANNs’ parameter estimates. Each panel shows the empirical distribution of the corresponding ANNs’ parameter estimate, for both groups (red: narrow range, blue: wide range). Panels are organized as follows: the left and right columns correspond to HP-ANN’s update rates and covariance thresholds, respectively ; the upper and lower rows correspond to ANNs with pseudo-gaussian and sigmoidal units’ activation functions, respectively.

Overall, empirical distributions of parameter estimates are qualitatively similar in both groups of participants. When comparing groups of participants with respect to either α_HP_ update rates, nothing reaches statistical significance (all p>0.4). This is also the case for HP covariance thresholds λ_HP_ under pseudo-gaussian activation functions (p=0.056, F=3.71, df=101). In addition, although HP covariance thresholds λ_HP_ under sigmoidal activation functions show a significant group difference (p=0.001, F=10.4, df=101), this difference is essentially driven by a few outlier individuals in the wide range group (cf. right tail of the distribution in the wide range group, lower-right panel of Figure S5).

### 7. ANNs’ behavioral postdiction and prediction accuracy

We fit a set of 2×2=4 ANNs (with and without hebbian plasticity, sigmoid versus gaussian activation functions) to each subject’s sequence of gamble decisions. In what follows, we provide summary statistics of our ANN-based behavioral.

Table S2 below gives the group average percentage of explained behavioral variance (R^2^) and its standard deviation (across participants) for each model (including the logistic regression model, for comparison purposes), and each group.

**Table S2:** Mean R^2^ and its standard deviation for each model, for both groups.

|  | narrow range group |  | wide range group |  |
| --- | --- | --- | --- | --- |
|  | mean | std | mean | std |
| logistic | 0.7789 | 0.1049 | 0.7367 | 0.1273 |
| ANN (gauss) | 0.8451 | 0.0784 | 0.8831 | 0.0866 |
| ANN (sig) | 0.8374 | 0.0793 | 0.7845 | 0.1166 |
| HP-ANN (gauss) | 0.8621 | 0.085 | 0.867 | 0.1673 |
| HP-ANN (sig) | 0.8527 | 0.0623 | 0.8225 | 0.127 |

Table S3 below gives the mean R^2^ difference between each ANN model and the reference logistic model, its standard deviation, and the resulting p-value (H0: no R^2^ difference), for both groups. Significant tests at the 5% level are highlighted in yellow.

**Table S3:** Mean R^2^ difference and its standard deviation for each ANN model, for both groups.

|  | narrow range group |  |  | wide range group |  |  |
| --- | --- | --- | --- | --- | --- | --- |
|  | mean | std | p-value | mean | std | p-value |
| ANN (gauss) | 0.0662 | 0.0939 | <10<sup>-4</sup> | 0.1463 | 0.0748 | <10<sup>-4</sup> |
| ANN (sig) | 0.0586 | 0.0446 | <10<sup>-4</sup> | 0.0477 | 0.0311 | <10<sup>-4</sup> |
| HP-ANN (gauss) | 0.0832 | 0.0601 | <10<sup>-4</sup> | 0.1302 | 0.167 | <10<sup>-4</sup> |
| HP-ANN (sig) | 0.0738 | 0.0841 | <10<sup>-4</sup> | 0.0857 | 0.119 | <10<sup>-4</sup> |

One can see that all ANNs achieve significantly better fit accuracy than the logistic model. This is expected however, since ANN models have much more degrees of freedom than the logistic model.

But is this improvement in fit accuracy homogeneous across decision difficulty? We then bin trials according to EV deciles, prior to averaging the rate of postdiction error across subjects. Figure S6 below summarizes the accuracy of behavioural postdictions as a function of gambles’ expected value, for all ANNs.

**Figure S6:**
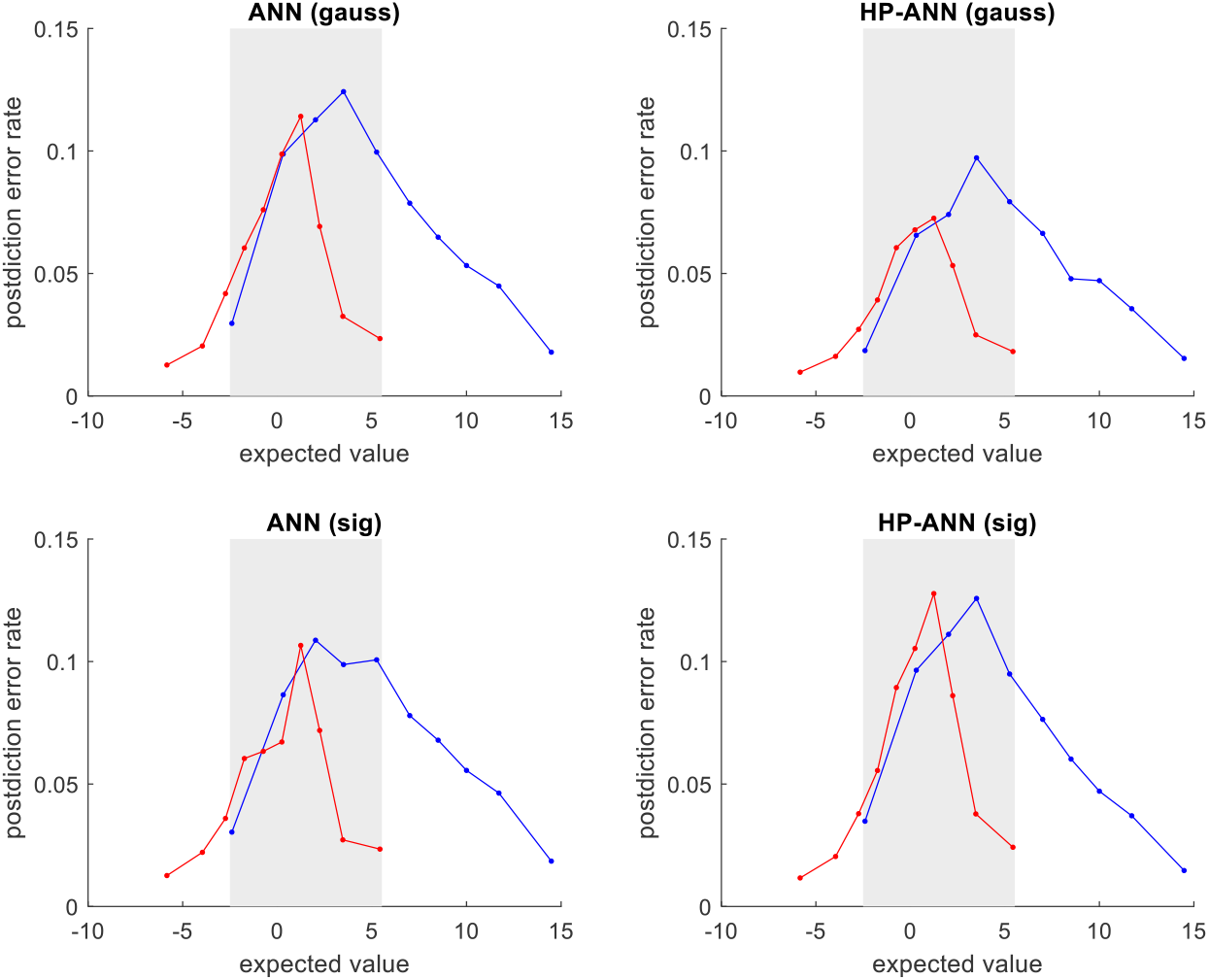
ANNs’ postdiction error rate. In each panel, the rate of postdiction error (y-axis) is plotted against gambles’ expected value (x-axis), for both groups (red: narrow range, blue: wide range). Same format as Figure S2. Panels are organized as follows: the left and right columns correspond to static ANNs and ANNs with Hebbian plasticity (HP-ANN), respectively ; the upper and lower rows correspond to sigmoid and pseudo-gaussian units’ activation functions, respectively.

As was the case for the logistic regression model, ANNs exhibit high fit accuracy for easy decisions (extreme EVs) and low fit accuracy for hard decisions (EV around 0). However, they yield much lower postdiction error rates, in particular for hard decisions (compare to Figure S2 above). In addition, sigmoidal default ANNs, pseudo-gaussian HP-ANNs seem to achieve better postdiction accuracy (below 11% error rate for hard decisions). But only HP-ANNs with gaussian activation functions generalize their predictions across risk range context (see Figure 3 in the main text).

### 8. ANN-RSA: assessing the statistical significance of model comparisons

Recall that our model space is factorial, with two orthogonal modelling factors: (i) our factor of interest has two “levels”: static ANN versus HP-ANN, and (ii) our factor of no interest also has two “levels”: sigmoidal versus pseudo-gaussian neural activation functions. This means that we will be comparing 2×2=4 models. When assessing the statistical significance of the ensuing model comparison, we will be using a variant of composite null testing.

Let 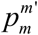 be the p-value associated with the elementary pairwise comparison of model *m* and *m*′, whose null hypothesis is 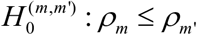, where *ρ_m_* is the corresponding Fisher-transformed RDM correlation (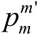 can be evaluated using paired t-tests on RDM correlations). Now for each model *m* ∈ [1,6], we ask whether its RDM correlation is the highest among candidate models. This induces the following composite null hypothesis: 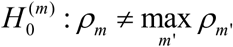. The maximum p-value statistics 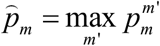 yields a valid test of the composite null hypothesis, though not necessarily maximally efficient (Wasserman, 2004). Because 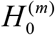 is the conjunction of elementary pairwise null hypotheses 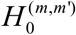, we refer to this approach as “conjunctive null testing”.

One may also want to evaluate the statistical significance of the comparison of RDM correlations across levels of our factor of interest, irrespective of our factor of no interest. The corresponding null hypothesis involves a disjunctive/conjunctive combination of elementary null hypotheses. For example, if one wants to test whether HP-ANNs exhibit a significantly higher RDM correlation than static ANNs (irrespective of the form of units’ activation functions), then the corresponding null hypothesis 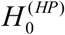 is defined as:

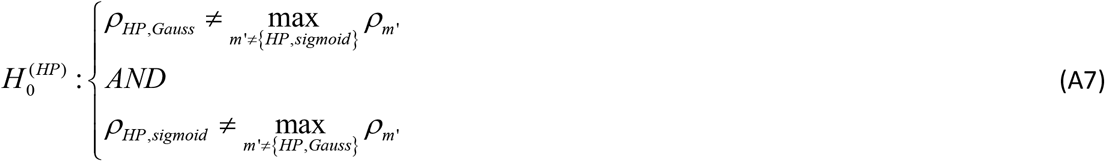

The following p-value then yields a valid statistical test of 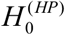:

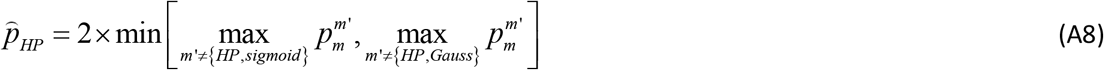

By design, the ensuing “disjunctive/conjunctive” approach cannot conclude about the underlying activation functions, i..e. it does not discriminate between sigmoid and pseudo-gaussian functional forms. However, it pools evidence over levels of our factor of no interest, which eventually improves statistical power. This is a frequentist -and simpler-variant of so-called “family inference” in Bayesian model comparison (Penny et al., 2010), where one marginalizes over modelling factors of no interest, effectively trading statistical power against inference resolution. We provide numerical evidence for the validity and efficiency of both conjunctive and disjunctive/conjunctive approaches below.

The critical question, for our combined ANN-RSA approach, it whether it exhibits the statistical robustness that is required for a reliable interpretation of fMRI data? For example, one may question the use of the parametric statistical approach above may not eventually result in inflated flase positive rates. But this also raises more basic modelling questions. In particular, one may ask whether the approach is robust to modelling assumptions regarding the (necessarily underestimated) dimensionality of ANNs that process behaviorally-relevant information. In other terms, does the approach accurately discriminate between candidate biological mechanisms of interest (e.g., ANNs with or without hebbian plasticity), when analyzing the data using ANN models that have much fewer activation units than the models that actually generate the data? We thus performed a series of Monte-Carlo simulations that recapitulates the design of the fMRI experiment.

We considered a decision task that requires the integration of two attributes *u* = {*G, L*} that vary randomly across 256 trials. We simulated four series of datasets, corresponding to the 2×2=4 ANN models. Each dataset was composed of 20 virtual subjects, whose trial-by-trial behavior and neural responses were generated under a “generative” ANN with sets of either *n_x_* = 20, 30 or 50 neural units. We introduced inter-individual variability by simulating dummy subjects with ANN parameters randomly sampled under their respective prior probability density functions (cf. Table S1 above). Each simulated multi-subject dataset was then analyzed using the ANN-RSA approach described above. In brief, each behavioral trial series was fitted with the 2×2 candidate ANNs, and the resulting estimated neural activity profiles were compared to simulated neural activity profiles using RSA. Importantly, fitted ANNs contained smaller sets of *n_x_* = 10 units than generative ANNs. For each dataset, we then compared models (at the group-level) using conjunctive and disjunctive/conjunctive null testing approaches. The former approach provides 4 p-values per dataset (one for each fitted ANN), while the latter provides 2 p-values per dataset (one for each ANN type, see above). We repeat this procedure 50 times, and keep track of all positive tests with a 5% significance threshold.

First, the conjunctive approach exhibits almost no model confusion. More precisely, the maximum false positive rate (about 10%) arises when testing static ANNs with gaussian activation functions, and only when the generative model was HP-ANN with gaussian activation functions and 30 units. This is likely due to random artefactual situations in which the generative ANNs exhibit very little plasticity. All other situations yield false positive rates much below 5%, which means that the test is valid. Second, the statistical power is variable: from about 92% ±2% on average for ANNs with sigmoid activation functions to about 31% ±25% on average for ANNs with gaussian activation functions. In other words, the conjunctive testing approach may be too conservative for ANNs with gaussian activation functions. Third, the dimensionality of generative ANNs seems to have almost no impact on statistical power. In other words, the conjunctive approach is robust to arbitrary modelling choices such as the number of ANN units.

Second, the confusion rate of positive disjunctive/conjunctive testing is similar to the conjunctive approach above. However, statistical power is much improved. Here again, the dimensionality of generative ANNs seems to have no impact on statistical power. Taken together, these results imply that, if a candidate mechanism eventually reaches statistical significance using either conjunctive or disjunctive/conjunctive approaches, then we can safely infer that it is a more likely explanation of fMRI activity patterns than other candidate mechanisms.

In conclusion, the ANN-RSA approach is robust to violations of modelling and statistical assumptions, including the low dimensionality of analyzing ANNs or the distribution of test statistics. We note that we expect this robustness to hold in more realistic situations, e.g., with network dimensionalities many orders of magnitude higher. This is because network units whose activation functions are centered far from the effective range of inputs either saturate or are silent, and thus do not contribute to the inter-trial covariation matrices (RDMs) that enter the RSA analysis. In fact, only those units whose activation function falls in between those of modelled units may eventually induce inter-trial covariations that would not be captured with small ANNs. However, this would only happen in situations where the behavioral response exhibit nonlinear effects that strongly deviate from smooth input integration profiles. This unlikely to be the case in our experimental context.

### 9. ANN-RSA analysis of fMRI data

Figure 4 of the Main Text only shows the results of the ANN-RSA analysis in OFC subregions. In addition to these three OFC subregions, 10 other ROIs were shown, in previous analyses of the NARPS dataset (Botvinik-Nezer et al., 2019, 2020), to respond to gains and/or to losses: namely, left/right striatum, left/right amygdala, left/right insula, left/right DLPFC, ACC and PCC (see Figure S7 below). In what follows, we report the results of the RSA analysis in all these ROIs.

**Figure S7:**
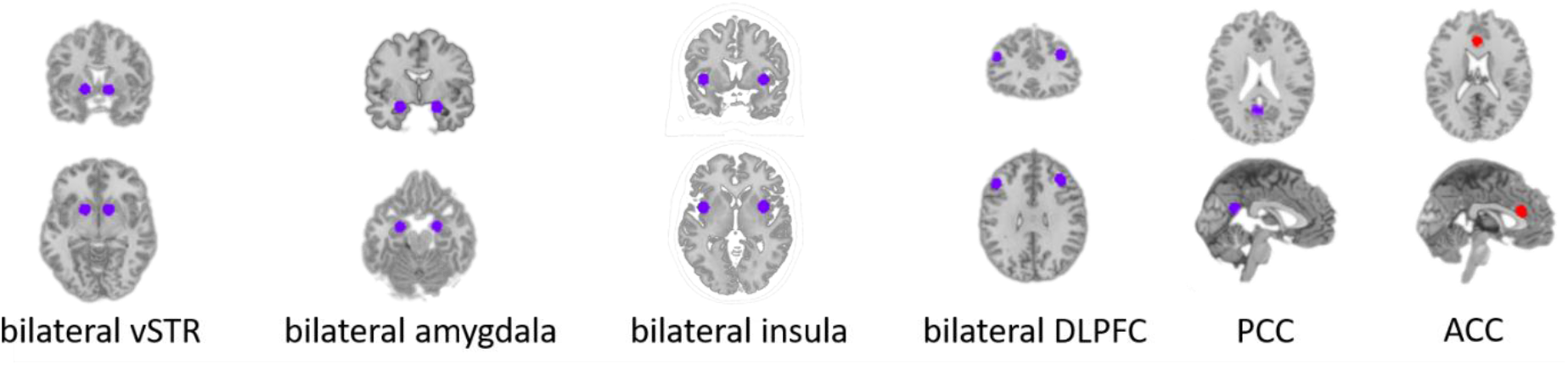
The location of each additional ROI is shown on the anatomical MRI template ; from left to right: left/right ventral striatum, left/right amygdala, left/right insula, left/right dorsolateral prefrontal cortex (DLPFC), posterior cingulate cortex (PCC) and anterior cingulate cortex (ACC).

To begin with, we simply ask whether either HP-ANNs (with pseudo-gaussian activation functions) actually explain multivariate fMRI time series, in any ROI that we included in our analysis. Figure S8 below summarizes this analysis, in terms of the group-average RDM correlations *ρ* for HP-ANNs and each ROI (and each group). Note that we correct all p-values for multiple comparisons across all 13 ROIs, including OFC subregions : for a target FPR at 5%, the Bonferroni-corrected significance threshold needs to be set to p=0.0038.

**Figure S8:**
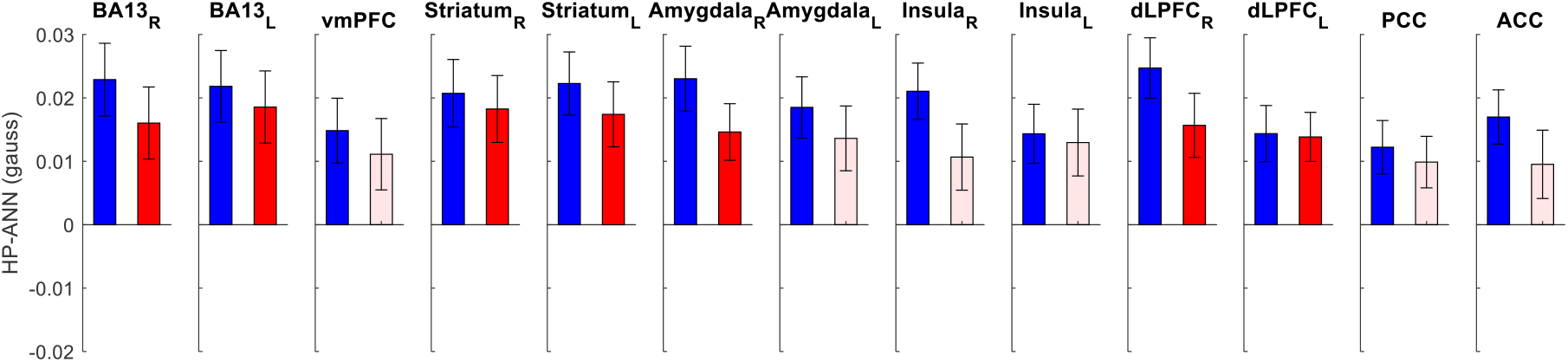
Each panel shows the group average RDM correlation ρ for HP-ANNs with pseudo-gaussian activation functions (blue: wide range group, red: narrow range group ; errorbars depict s.e.m), for each ROI. Plain bars depict statistical significance (at FPR=5%, with Bonferroni correction across all ROIs), whereas shaded bars show non significant results.

Most importantly, RDM correlations in bilateral OFC (left and right BA13) remain significant in both groups of participants, even when correcting for multiple comparisons across all ROIs.

Now let us summarize the RSA results in the additional ROIs. In brief, HP-ANNs are the only candidate models that reach statistical significance (all other ANNs: p>0.04 uncorrected). Furthermore, the only ROIs where HP-ANNs yield significant RDM correlations *in both groups* are the bilateral ventral striatum, bilateral DLPFC, and right amygdala. We then compared Hebbian plasticity to other biological mechanisms of interest using disjunctive/conjunctive testing (see methodological section above). In the narrow range group, we found that HP-ANNs exhibit significantly higher RDM correlations than other ANNs in bilateral Striatum (left Striatum: p=0.0003, right Striatum: p=0.001) and in the right Amygdala (p=0.0006). In other ROIs, no comparison of RDM correlations achieves statistical significance (all p>0.1). Furthermore, the RDM correlations of static ANNs are never significantly higher than those of other ANNs (all p>0.095, uncorrected). The results of the wide range group remarkably replicate those of the narrow range group: the RDM correlations of HP-ANNs are only significantly higher than other ANNs in bilateral Striatum (left Striatum: p=0.0018, right Striatum: p=0.0006), although there is a trend in bilateral Amygdala (left/right Amygdala: p=0.0076). Finally, the RDM correlations of static ANNs are nowhere statistically higher than those of other ANNs.

For completeness, Tables S4 (resp. S5) below gives the p-value of RDM correlations for each model and each ROI (H_0_: *ρ* ≥ 0, one-sided t-test), in the narrow (resp. wide) range group. The tests that are significant at FWE=5% are highlighted in yellow.

**Table S4:**
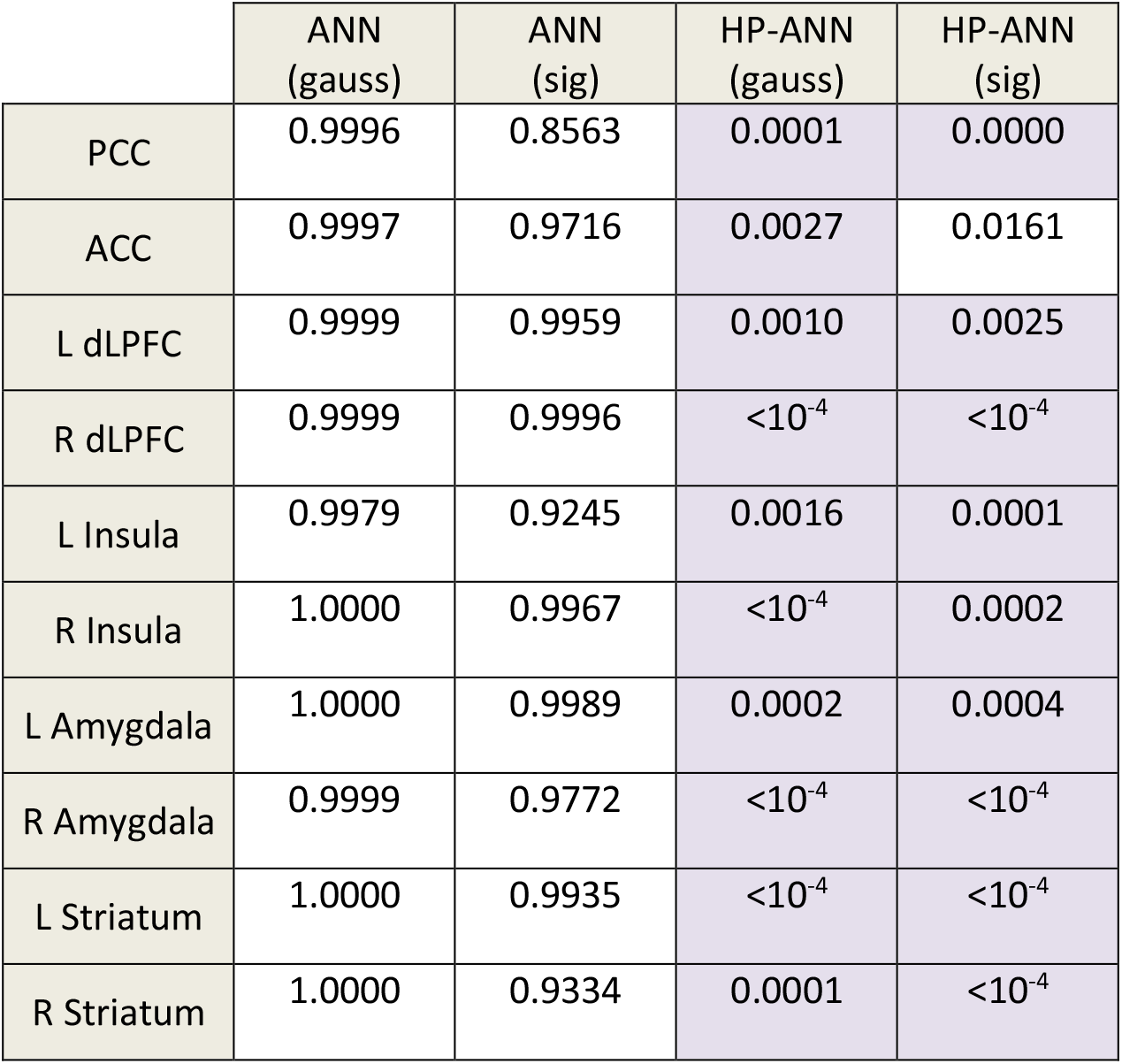
P-value of RDM correlations for each model and each ROI (wide range group).

|  | ANN<br>(gauss) | ANN<br>(sig) | HP-ANN<br>(gauss) | HP-ANN<br>(sig) |
| --- | --- | --- | --- | --- |
| PCC | 0.9996 | 0.8563 | 0.0001 | 0.0000 |
| ACC | 0.9997 | 0.9716 | 0.0027 | 0.0161 |
| L dLPFC | 0.9999 | 0.9959 | 0.0010 | 0.0025 |
| R dLPFC | 0.9999 | 0.9996 | $<10^{-4}$ | $<10^{-4}$ |
| L Insula | 0.9979 | 0.9245 | 0.0016 | 0.0001 |
| R Insula | 1.0000 | 0.9967 | $<10^{-4}$ | 0.0002 |
| L Amygdala | 1.0000 | 0.9989 | 0.0002 | 0.0004 |
| R Amygdala | 0.9999 | 0.9772 | $<10^{-4}$ | $<10^{-4}$ |
| L Striatum | 1.0000 | 0.9935 | $<10^{-4}$ | $<10^{-4}$ |
| R Striatum | 1.0000 | 0.9334 | 0.0001 | $<10^{-4}$ |

**Table S5:** P-value of RDM correlations for each model and each ROI (narrow range group).

|  | ANN<br>(gauss) | ANN<br>(sig) | HP-ANN<br>(gauss) | HP-ANN<br>(sig) |
| --- | --- | --- | --- | --- |
| PCC | 0.1574 | 0.7610 | 0.0418 | 0.0059 |
| ACC | 0.2220 | 0.9285 | 0.0094 | 0.0674 |
| L dLPFC | 0.0375 | 0.9935 | 0.0004 | 0.0575 |
| R dLPFC | 0.0398 | 0.5038 | 0.0016 | 0.0009 |
| L Insula | 0.2706 | 0.6680 | 0.0088 | 0.0192 |
| R Insula | 0.4798 | 0.5675 | 0.0233 | 0.0031 |
| L Amygdala | 0.6118 | 0.7747 | 0.0052 | 0.0006 |
| R Amygdala | 0.1554 | 0.4109 | 0.0010 | 0.0005 |
| L Striatum | 0.2719 | 0.1865 | 0.0007 | 0.0002 |
| R Striatum | 0.1803 | 0.2060 | 0.0006 | 0.0007 |

One can see that the shape of the units’ activation functions (either pseudo-gaussian or sigmoidal) has virtually no impact on RSA results.

### 10. On the diversity of response profiles in HP-ANNs’ integration units

OFC neurons are notoriously diverse in their response profile, but a consistent finding is that, in the context of value-based decision making, they can be classified in terms of so-called ‘choice cells’, ‘chosen value cells’ and ‘offer value cells’ (Padoa-Schioppa and Assad, 2006, 2008). Given that this can be considered a pre-requisite for any computational model of value integration in the OFC, we asked whether HP-ANNs re produce this known property of OFC neurons.

For each subject, we thus tested whether the response of integration units correlates (across trials) with choice, chosen value and/or gamble value, where value is defined as the weighted sum of gains and losses (according to the static logistic model parameter estimates). Integration units are then classified in terms of which variable it correlates most with (but units are attributed none of these labels if the correlation is not significant). The results of this analysis are summarized on Figure S9 below.

**Figure S9:**
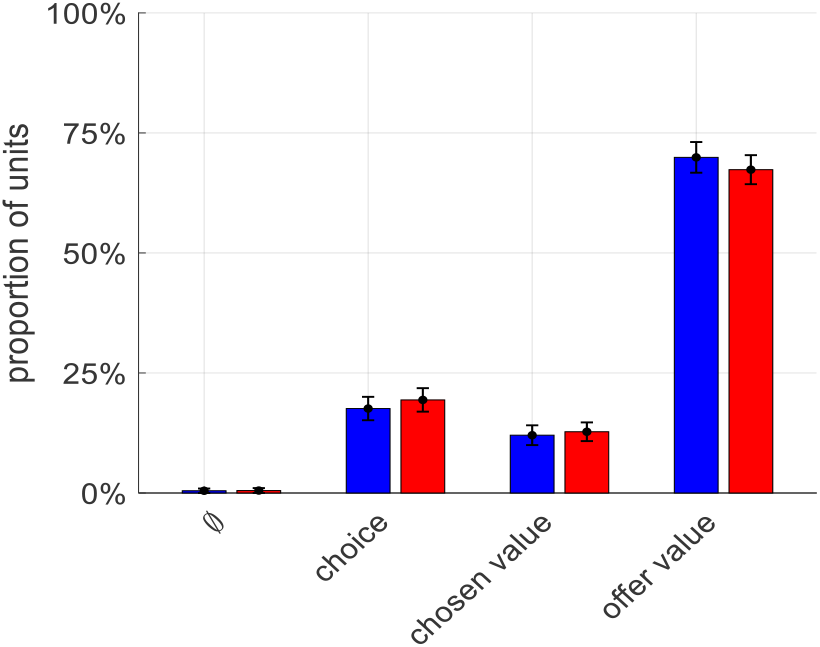
response profile diversity in HP-ANNs’ integration units. The average proportion of HP-ANNs integration units (y-data) that shows a significant correlation with choice, chosen value, and offer value (or none of these) is plotted for for both groups (blue: wide range, red: narrow range). Errorbars depict s.e.m. (across participants).

We find that, although HP-ANN’s integration units were not at all designed to encode these quantities, they eventually reproduce the response variability observed in OFC neurons. Importantly, the response profiles are almost identical for both groups. In particular, we find that about 13% of integration units are classified as “chosen value” cells. This is interesting, because the computational role of these units is in fact exactly the same as that of units that are classified as “offer value” cells: together, they form a population code for the subjective value of gambling. In other words, this classification is not directly relevant for guessing the underlying computational role of integration units.

